# A retinotopic wiring principle of the human brain

**DOI:** 10.64898/2026.04.03.716412

**Authors:** G. Amorosino, B. Caron, J. Kwon, M. Carrasco, R.C. Reid, C. Lenglet, J. Zimmermann, E. Yacoub, K. Ugurbil, S.R. Heilbronner, F. Pestilli

## Abstract

A central challenge in neuroscience is to understand how anatomical connectivity links neural representations to behavior. In sensory systems, cortical areas encode structured maps of the external world, yet it remains unclear whether and how this representational geometry constrains long-range connectivity in the living human brain. Here, we show that structural connectivity in the human visual cortex preserves retinotopic organization, revealing a retinotopic wiring principle; cortical regions representing the same locations in visual space preferentially connect. Across more than 1,700 participants spanning ages 2 to 88 years, we further show that well-known perceptual asymmetries correspond to systematic asymmetries in connectivity within early visual cortex, but not in connections linking visual cortex to the rest of the brain. We introduce a scalable approach to map retinotopic connectivity when tractography alone is unreliable. Together, these findings demonstrate that anatomical connectivity preserves the geometry of neural representations, providing a general principle linking brain structure, function, and behavior.

## Main

Visual perception emerges from coordinated activity across multiple cortical areas that represent the same visual scene. Although decades of research have revealed the functional organization of these areas and the response properties of the neurons they contain^1–9^ significantly less is known about how they interact through structural connections to form an integrated visual network. In particular, the set of white-matter pathways linking retinotopic visual maps across cortex—the visual connectome—remains incompletely characterized in the human brain. As a result, although the components of the visual system are well defined, the structural architecture that coordinates activity across visual maps to support perception remains elusive.

A defining property of early visual cortex is retinotopy: the spatial organization in which neuronal responses depend on stimulus position in the visual field^10^. Neighboring cortical neurons represent neighboring locations in visual space, producing maps that preserve the geometry of the visual environment (**Fig. 1a**). Early visual areas—including V1, V2, V3, hV4, VO1, VO2, LO1, LO2, TO1, TO2, V3a, and V3b—encode features such as orientation, spatial frequency, color, and contrast, with representations becoming progressively more invariant along the visual hierarchy^11^. Because multiple visual areas contain aligned retinotopic maps, perception depends on coordinated interactions across cortical locations representing corresponding positions in visual space. This raises the possibility that structural connectivity is organized in the coordinate system of visual space: A retinotopic connectivity principle. We performed two tests to establish the retinotopic wiring principle.

**Figure 1.**
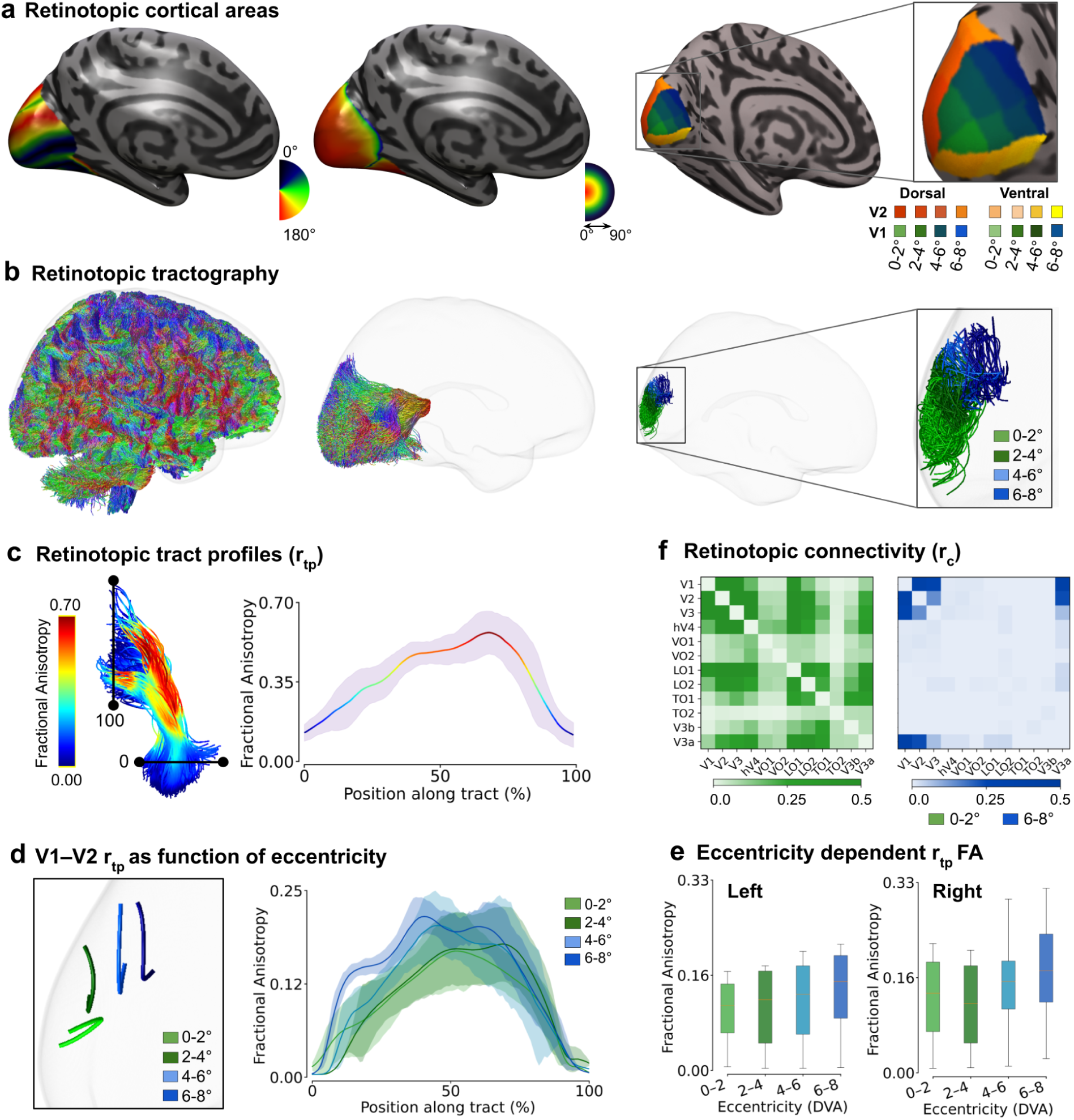
Retinotopic connectivity and white matter. **a.** Schematic illustration of a retinotopic cortical organization. *Left*, posterior views of the inflated cortical surface illustrating retinotopic organization of the visual cortex. The first panel shows the polar angle map, and the second panel shows the eccentricity map. *Right*, schematic representation of polar angle and eccentricity coordinates in the visual field. **b.** Retinotopic tractography linking corresponding visual field representations between V1 and V2. **c.** *Retinotopic tract profiles*, *r_tp_*, reconstructed white-matter tract connecting retinotopic V1 and V2 regions. *Right*, retinotopic tract profile showing microstructural measurements along the pathway. **d.** Representative retinotopic tract profiles variations in fractional anisotropy (FA) as a function of eccentricity (mean ±standard error (s.e.) across the tract). **e.** Retinotopic FA increases as a function of eccentricity in both the left and right hemisphere (n=1, mean ±standard error (s.e.) across the tract). **f.** *Retinotopic connectivity*, *r_c_*, for pairs of twelve visual areas (V1, V2, V3, V3A, V3B, hV4, LO1, LO2, TO1, TO2, VO1, and VO2). Colormap, reproduces *S_d_*, Eq. 1.

Insights from cellular and mesoscale connectomics suggest a potential organizing principle. In mouse visual cortex, neurons preferentially connect to other neurons with similar functional properties, a phenomenon often described as *like-to-like* connectivity^12,13^. If similar wiring rules operate across cortical areas in the human brain, then cortical locations representing the same position in visual space should preferentially connect across retinotopically aligned maps. Testing this prediction in the living human brain, however, has remained technically challenging.

Retinotopic organization also shapes several well-established perceptual phenomena^14–18^. Visual performance varies systematically across the visual field: observers typically show superior performance along the horizontal relative to the vertical meridian (the horizontal-vertical asymmetry, HVA), and along the lower relative to the upper visual field (the vertical meridian asymmetry, VMA;^19^. These perceptual asymmetries correspond to differences observed in cone and midget retinal ganglion cell densities^20,21^, which are exacerbated by cortical magnification and functional responses across early visual cortex^22–24^. Such findings suggest that the geometry of visual perception could reflect the underlying architecture of the visual system. However, whether these perceptual asymmetries arise in part from differences in structural connectivity between visual maps remains unknown.

Diffusion-weighted magnetic resonance imaging combined with tractography provides the only non-invasive method for mapping white-matter pathways in vivo. However, tractography reconstructs anatomical pathways without preserving the visual field coordinates that define cortical maps, limiting the study of connectivity between homologous retinotopic locations. As a result, most studies characterize connectivity between coarse anatomical regions^25^ rather than within the coordinate system of visual space, and retinotopically specific connectivity has remained difficult to measure beyond a small number of well-characterized pathways, e.g., the optic radiation^26–28^, the vertical occipital fasciculus^29,30^, or small sets of V1–V2 connections obtained using highly specialized hardware that is not widely accessible, constraining scalability to large populations^31–33^).

Here we introduce an automated framework for mapping retinotopically specific structural connectivity across the human visual system. Applying this framework to more than 1,700 participants aged 2–88 years across multiple datasets, we reconstruct visual pathways as a function of eccentricity and polar angle. We show that connectivity between visual areas follows a retinotopic wiring principle, whereby cortical regions representing the same locations in visual space preferentially connect. We further demonstrate that well-known perceptual asymmetries correspond to systematic asymmetries in connectivity within early visual cortex, but not in connections linking visual cortex to the rest of the brain. Finally, we introduce a scalable approach for constructing population-level retinotopic connectivity templates, enabling robust inference of visual pathways when tractography alone is unreliable. Together, these results demonstrate that structural connectivity preserves the geometry of neural representations, providing a framework for linking brain structure, function, and behavior.

## Results

Structural connectivity in the human visual cortex preserves retinotopic organization, such that cortical regions representing similar locations in visual space preferentially connect across areas. To test this, we developed VISCONN (VISion CONNectivity toolbox; https://visconn.github.io/), a framework that integrates retinotopic maps with diffusion tractography to reconstruct white-matter pathways as a function of visual field position defined by eccentricity and polar angle (**Fig. 1**, see also **Supplementary Fig. 1** and **2**). VISCONN enables estimation of connectivity between cortical locations representing the same position in visual space across visual areas, overcoming a central limitation of conventional tractography, which does not preserve retinotopic coordinates. VISCONN builds on established software libraries for anatomical and tractography processing^34–39^, together with anatomical models of retinotopic organization in early visual cortex^40,41^. The functionality of these tools was customized and optimized to segment twelve retinotopic areas—V1, V2, V3, V3A, V3B, hV4, LO1, LO2, TO1, TO2, VO1, and VO2—and parcellate each area according to its representation of visual field coordinates defined by eccentricity and polar angle (**Fig. 1a**; see^40,41^). These cortical segmentations were combined with tractography outputs to reconstruct connections between retinotopic locations across visual maps. The resulting connectivity estimates are summarized as retinotopic connectivity matrices and tract profiles, both indexed by visual field position (**Fig. 1b**).

**Figure 2.**
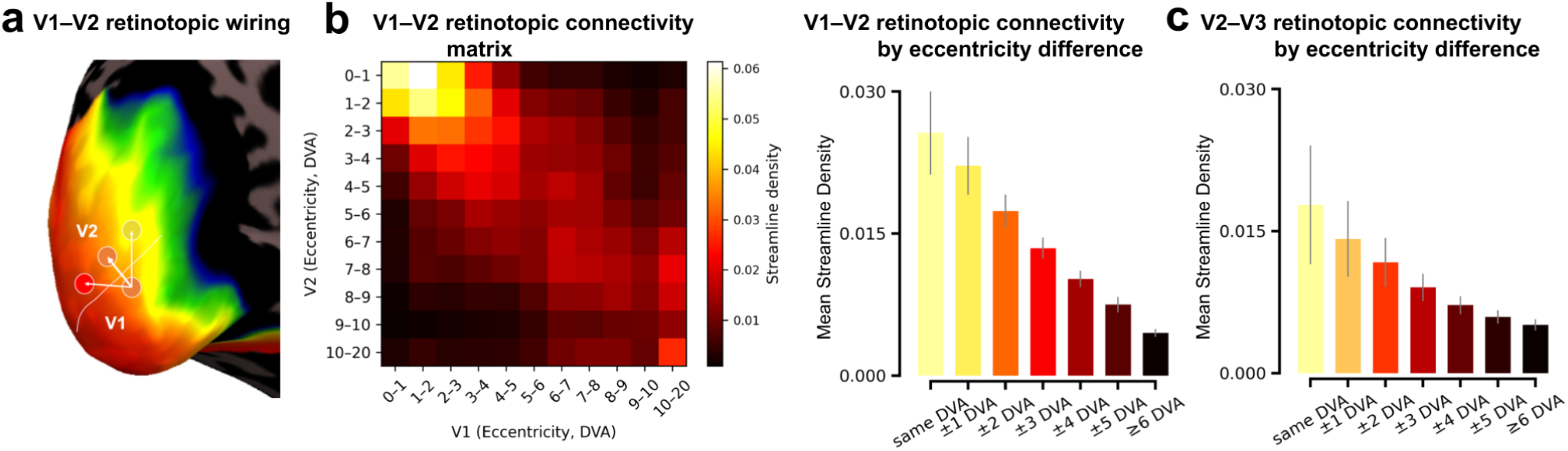
Like-to-like retinotopic connectivity across visual areas. **a**, Schematic of retinotopically defined eccentricity segments in V1 and their connections to corresponding segments in V2. **b**, Connectivity between V1 and V2 as a function of eccentricity. *Left*, eccentricity-by-eccentricity matrix of streamline density (S_d_) showing strongest connectivity along the diagonal, indicating preferential connections between homologous visual field locations. *Right*, mean S_d_ as a function of eccentricity distance, demonstrating maximal connectivity between matching eccentricity bins (same degrees of visual angle, DVA) and a monotonic decline with increasing separation. **c**, Retinotopic connectivity between V2 and V3 shows a similar but weaker organization, indicating that like-to-like retinotopic connectivity is preserved across visual areas while decreasing in strength along the visual hierarchy All values represent mean ± s.e.m.

We analyzed diffusion MRI data from 1,061 participants from the Human Connectome Project (HCP)^42^. In each participant, continuous polar angle and eccentricity maps were first estimated for V1 and V2 using automated anatomical-prediction methods^40,41^ and each map was then subdivided into four eccentricity bins (*b_e_*, 0-2°, 2-4°, 4-6° and 6-8° DVA) and each bin was further partitioned into ventral and dorsal sectors corresponding to lower and upper visual field representations, respectively (ventral bins, 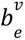 0-90 polar angle, and dorsal bins, 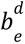 90-180° polar angle; **Fig. 1a**). Whole-brain anatomically constrained probabilistic tractography was then performed to generate approximately three million streamlines per brain^43^. Retinotopic connections between visual maps were identified by assigning streamline endpoints to the gray–white matter interface of aligned retinotopic locations across cortical areas (**Fig. 1b**).

Retinotopic tractography was summarized through two complementary representations. First, the microstructural properties of pathways linking specific visual field locations can be characterized using tract profilometry, which we refer to as *retinotopic tract profiles* (*r_tp_*). As an illustration, we estimated fractional anisotropy (FA) along dorsal V1–V2 for each *b_e_* (**Fig. 1c**). Across the cohort, mean FA increased systematically with eccentricity, consistent with behavioral reports indicating faster processing speeds in the visual periphery (**Fig. 1d,e**)^44–47^. Second, VISCONN quantifies *retinotopic connectivity* (*r_c_*) as the matrix of structural connectivity estimated between matched retinotopic positions across pairs of visual areas. We estimated *r_c_* among twelve early visual areas (V1, V2, V3, V3A, V3B, hV4, LO1, LO2, TO1, TO2, VO1, and VO2; **Fig. 1f**) using streamline density (*S_d_*; *Eq. 1*, **Methods**) across eccentricity bins. Overall, visual areas’ *r_c_* decreased as function of eccentricity (see also **Supplementary Fig 3**). Although the decrease *r_c_* across eccentricity may appear opposite the reported V1-V2 *r_tp_* trend, the result is consistent with information integration and receptive field size increase in peripheral vision^11^. The VISCONN toolbox is distributed through brainlife.io^48^ as a reproducible collection of ready-to-use applications (**Supplementary Table 1**). In the following sections, we test whether structural connectivity between visual maps follows a fundamental retinotopic organizational principle in two ways. First, examine whether connectivity between visual areas follows a retinotopically specific “*like-to-like*” wiring principle: cortical locations representing the same position in visual space preferentially connect across areas.

### Like-to-like connectivity in the human visual system

Recent connectomics studies in mice have shown that neurons with similar functional properties preferentially form synaptic connections, a phenomenon termed the *like-to-like* principle^12,13^. In these studies, like-to-like organization was defined at the cellular scale using orientation selectivity and synaptic connectivity. Here, we tested whether an analogous organizational principle is present in the human visual system at the mesoscale (millimeter scale). In contrast to the murine studies, we examined long-range axonal connectivity between visual cortical areas using diffusion MRI tractography constrained by retinotopic organization. Specifically, we evaluated whether regions representing similar visual field positions preferentially connect across areas of human visual cortex. Analyses were conducted using diffusion MRI data from 1,061 participants from the Human Connectome Project (HCP). Specifically, we tested whether iso-eccentric locations in pairs of early visual areas preferentially connect. Because orientation-selective synaptic connectivity cannot currently be measured *in vivo* in the human brain, our analysis focused on a spatial generalization of these rodent findings. In this formulation, the *like-to-like* principle predicts that cortical locations at the *mm* scale, representing the same position in visual space, will preferentially connect across retinotopically aligned visual maps.

To test this prediction, we focused on two canonical pairs of visual areas, V1–V2 and V2–V3, and subdivided each region into 1° eccentricity bins (*b_e_*, **Fig. 2a** *left*), spanning from 0° to 20° DVA (**Fig. 2a**, *left*). Structural connectivity was then quantified between all possible *b_e_*-pairs across the two areas. Specifically, streamline density (*S_d_*) was estimated between each pair of V1–V2 *b_e_*, producing eccentricity-by-eccentricity retinotopic connectivity matrices (*r_c_*, **Fig. 2a** *right*). Under a like-to-like principle, structural connectivity is predicted to be strongest between iso-eccentric locations, resulting in a prominent diagonal structure within the *r_c_*matrices.

The *like-to-like* principle predicts that iso-eccentric *b_e_*(main-diagonal elements of *r_c_*) should exhibit higher *S_d_*than non-iso-eccentric *b_e_* (off-diagonal *r_c_* elements). To quantify this effect, we averaged *r_c_* values along successive off-diagonals representing increasing eccentricity differences between *b_e_*pairs. Across 1,061 participants drawn from HCP datasets, the results revealed a robust *like-to-like* retinotopic pattern of connectivity. For V1–V2, connectivity was strongest along the main diagonal of the *r_c_* matrix (**Fig. 2b**, *left*; χ²(6) = 5370.98, p < 0.001, Friedman’s ANOVA; see **Supplementary Table 2**). Connectivity declined sharply beyond ±3° DVA (p < 0.001, Holm-corrected, Wilcoxon signed-rank; see **Supplementary Table 3**). A similar organization was observed for V2–V3 connections (χ²(6) = 4510.36, p < 0.001, Friedman’s ANOVA; see **Supplementary Table 2**), with connectivity peaking at iso-eccentric locations and decreasing thereafter (**Fig. 2b**, *right*). Connectivity declined significantly less markedly beyond ±3° DVA (p < 0.001, Holm-corrected, Wilcoxon signed-rank; see **Supplementary Table 4**). The shallower *like-to-like* decline in V2–V3 is consistent with the known expansion of receptive field sizes along the vertical hierarchy, which leads to greater integration of information across the visual field^11^. To account for potential distance-related biases in tractography^49^, we normalized streamline density (*S_d_*) by the mean streamline length for each *b_e_* pair (S̅_*d*_; **Supplementary Results 3**). The like-to-like connectivity pattern was reduced in magnitude, but persisted after normalization, demonstrating that the effect is not explained by differences in inter-regional distance (**Supplementary Fig. 4**). Together, these results indicate that structural connectivity between visual areas follows a retinotopically specific *like-to-like* organization, linking cortical locations with homologous visual space representation.

### Retinotopic connectivity corresponds to perceptual asymmetries

Having established that a fundamental wiring principle of the rodent visual system is conserved in the human brain, we next asked whether these structural connections correspond to well-documented asymmetries in human perception. At iso-eccentric locations, human adults exhibit higher perceptual performance along the horizontal than vertical meridian (HVA; Eq. 2). Perceptual performance is also higher along the lower than the upper vertical meridian (VMA; *Eq. 3*). Both the HVA and VMA have been consistently observed across a wide range of visual tasks, including contrast sensitivity^50–59^, spatial resolution and acuity^60–69^, motion perception^70–72^, and short-term memory^64^. Previous work has linked these asymmetries to structural and functional properties of the visual system, including variations in cortical surface area within early visual cortex—particularly V1^15,24,73–80^—and differences in functional imaging responses^23^ measured with fMRI (see ref^19^ for a review). However, whether the retinotopic connectivity between visual areas relates to these perceptual asymmetries remains largely unexplored.

To test this hypothesis, we analyzed diffusion MRI data from over 1,700 participants spanning three independent datasets and different age groups: 1,061 participants from the HCP^42^, 106 from the Pediatric Imaging, Neurocognition, and Genetics study (PING)^81^, and 584 from the Cambridge Centre for Ageing and Neuroscience (Cam-CAN)^82^. These cohorts span multiple acquisition protocols and imaging resolutions, collectively covering most of the human lifespan (ages, 2 to 88). Using VISCONN, we segmented the cortical surfaces of V1 and V2 into three regions of interest (ROIs) defined by polar meridian and eccentricity: the upper vertical meridian (UVM; eccentricity 0–90°; polar angle: ±15°), the lower vertical meridian (LVM; eccentricity 0–90°; polar angle: 165–195°), and the horizontal meridian (HM; eccentricity 0–90°; polar angle: 75–105°). The vertical meridian (VM) was defined as the union of the UVM and LVM (**Fig. 3a** *left*).

**Figure 3.**
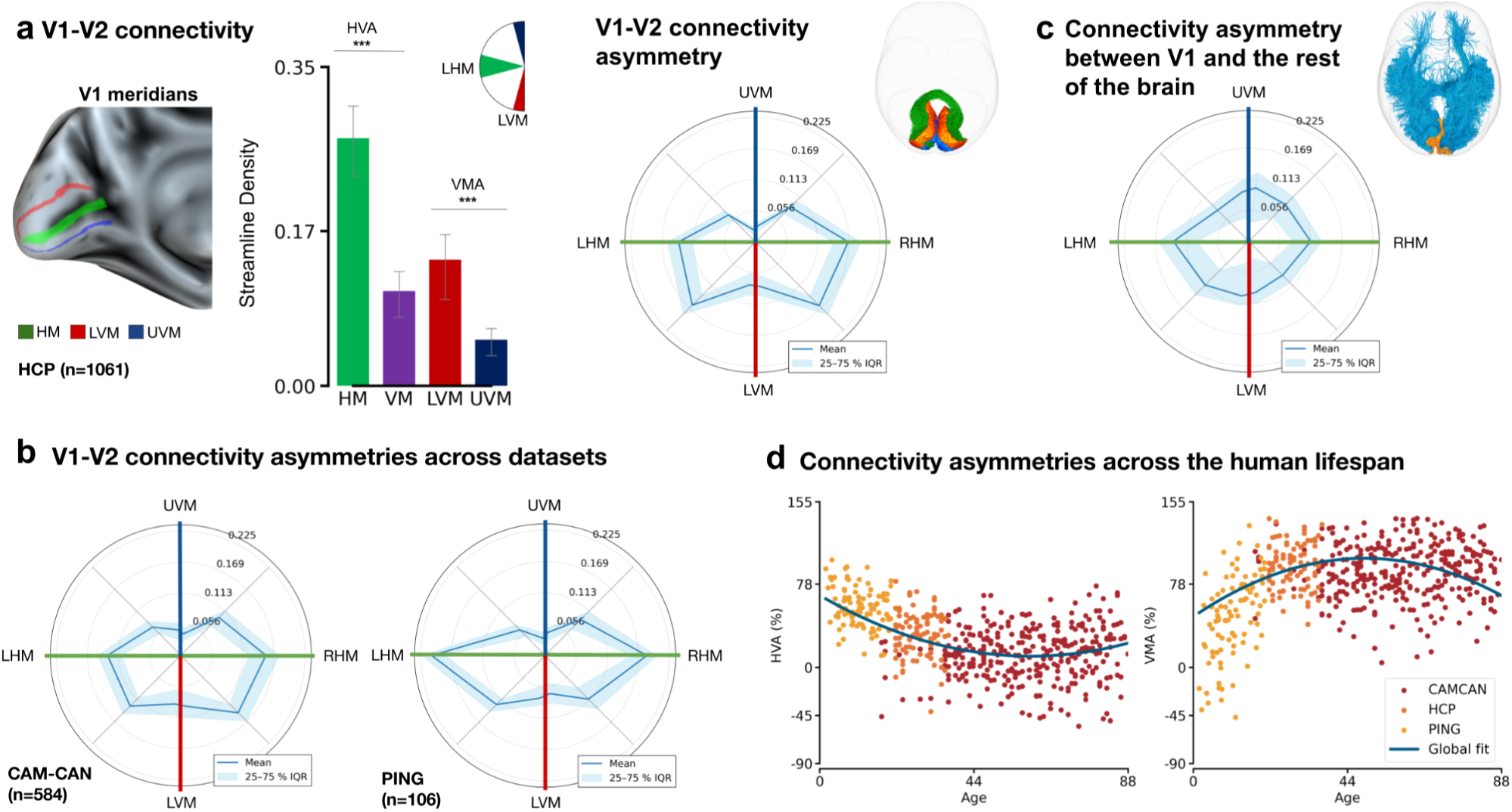
Visual connectivity asymmetries. **a.** Meridian asymmetries in early visual white-matter connectivity. *Left*, surface-based representations of the horizontal meridian (HM) and vertical meridian subdivisions, including the lower vertical meridian (LVM) and upper vertical meridian (UVM), in V1. *Middle*, bar plots showing mean streamline density (*S_d_*) across the three principal meridians (HM, LVM, and UVM). *Right*, polar-angle–resolved connectivity profiles between V1 and V2, demonstrating higher connectivity along the horizontal compared with the vertical meridian and along the lower compared with the upper vertical meridian. Results were derived from diffusion MRI tractography using the Human Connectome Project (HCP) dataset (n = 1,061 participants). **b.** Replication across independent cohorts. Meridian asymmetry profiles derived from the Cambridge Centre for Ageing and Neuroscience (Cam-CAN, n = 584) and Pediatric Imaging, Neurocognition, and Genetics (PING, n = 106) datasets reproduce the horizontal–vertical asymmetry (HVA) and vertical meridian asymmetry (VMA) patterns. **c.** Meridian asymmetries are absent in long-range visual connectivity. Polar-angle–resolved streamline density profiles for connections between the rest of the brain and V1 show reduced or absent asymmetries, indicating that meridian-dependent connectivity differences arise primarily within the visual system. **d.** Lifespan trajectories of meridian asymmetries. Individual HVA (left) and VMA (right) indices plotted as a function of age across PING, HCP, and Cam-CAN reveal distinct developmental and aging-related trends. For visualization purposes, data points were histogram-matched across datasets.

Structural connectivity was quantified as streamline density (*S_d_*) between corresponding ROIs across visual areas. Connectivity was highest between HM representations in V1 and V2, lowest between UVM representations, and intermediate between LVM representations (**Fig. 3a**, *left*). When visualized as a function of position in the visual field, these connectivity patterns closely resembled the spatial distribution of perceptual performance across the visual field (**Fig. 3a**, *right*; see^19^). A significant effect of visual meridian on *S_d_*was observed (p < 0.001, Friedman’s ANOVA; see **Supplementary Table 5**), indicating systematic differences across HM, LVM, and UVM representations. These effects were consistently observed across independent datasets (**Fig. 3b**), demonstrating that the overall pattern of connectivity asymmetries generalizes across imaging protocols and participant populations. Results further revealed a robust vertical meridian asymmetry (VMA; LVM > UVM; p < 0.001,

Holm-corrected, Wilcoxon signed-rank; see **Supplementary Table 6**), which was present across all datasets (HCP: Cohen’s d = 1.47; Cam-CAN: d = 0.96; PING: d = 0.8).A significant horizontal–vertical asymmetry was also present in all datasets (HVA; HM > VM; all p < 0.0001), with large effect sizes in HCP (d=2.51), Cam-CAN (d=1.38), and PING (d= 2.32). These asymmetries were also observed for V1–V3 and V1–hV4 connections (V1–V3: χ^2^(2) = 1943.81, p < 0.001; V1–hV4: χ^2^(2) = 2011.75, p < 0.001; **Supplementary Fig. 5**), but in later visual areas only the the VMA, but not the HVA is present (**Supplementary Fig. 6**), suggesting that retinotopic connectivity asymmetries might arise from early visual areas. Because diffusion tractography is susceptible to gyral bias (favoring connections between gyral crowns over sulcal walls^83^) we performed control analyses to determine whether this bias could account for the observed connectivity asymmetries. The results show that the asymmetries persist after accounting for gyral bias (**Supplementary Results 6**).

We next asked whether the HVA and VMA arise primarily from local connectivity within the visual system or whether similar asymmetries are also present in connections between visual cortex and the rest of the brain. To address this question, we estimated long-range connections between V1 meridian ROIs and all cortical regions outside the occipital lobe (excluding the twelve visual areas; **Fig. 3c** and **Methods**). In contrast to the strong meridian-dependent effects observed within visual cortex, global connections did not exhibit significant HVA or VMA patterns (HVA: d = −0.63, p = 1.0; VMA: d = 0.02, p = 0.41). These findings indicate that perceptual asymmetries are specifically reflected in the organization of connectivity within the visual network, rather than in the large-scale connections linking the visual cortex and the broader brain.

Finally, we examined how connectivity asymmetries vary across the human lifespan (ages 2–88). Previous studies investigating structural and behavioral asymmetries in vision have reported age-related changes, from childhood to adulthood, particularly within the vertical meridian asymmetry^84,85^. To evaluate this relationship, we fit a quadratic model to subject-specific estimates of HVA and VMA across the combined datasets (**Fig. 3d**, *Eq. 4*). Whereas the HVA showed only a positive quadratic relationship with age (R² = 0.12; AIC = 11121), the VMA exhibited a negative quadratic relationship with age (R² = 0.09; AIC = 11300). These results indicate that structural connectivity asymmetries vary across the lifespan suggesting that the vertical meridian asymmetry is particularly sensitive to developmental and aging processes. Intriguingly, the perceptual VMA continues to mature throughout adolescence and only reaches an adult state by late adolescence^85^—a period in which the connectivity VMA is also increasing. More generally, these results demonstrate a fine match between established perceptual phenomena and the retinotopic connectivity principle.

### Population templates recover retinotopic connectivity beyond tractography limitations

Population-level templates recover retinotopically specific connectivity patterns that are not reliably detected in individual tractography. Although our approach reconstructs visual pathways at the group level (**Fig. 1-3**), reconstructions in individual subjects vary in completeness, consistent with known limitations of diffusion tractography^86,87^. For example, connections between specific visual areas may be partially or inconsistently reconstructed across individuals.

To address this limitation, we developed a population-based approach that aggregates tractography across participants to estimate a consensus representation of retinotopic connectivity. By combining occipital streamlines from more than 1,000 participants from the HCP dataset, we constructed a compact population template (*T_t_*) that captures the stable geometric organization of visual pathways while reducing noise and reconstruction failures at the individual level (**Fig. 4a**; see **Methods**). Results show that *T_t_* preserves the geometric organization of visual pathways while remaining computationally efficient. *T_t_* provides a compact population-level reference for retinotopic connectivity across visual areas and enables inference of visual pathways in individual brains even when tractography is unreliable.

**Figure 4.**
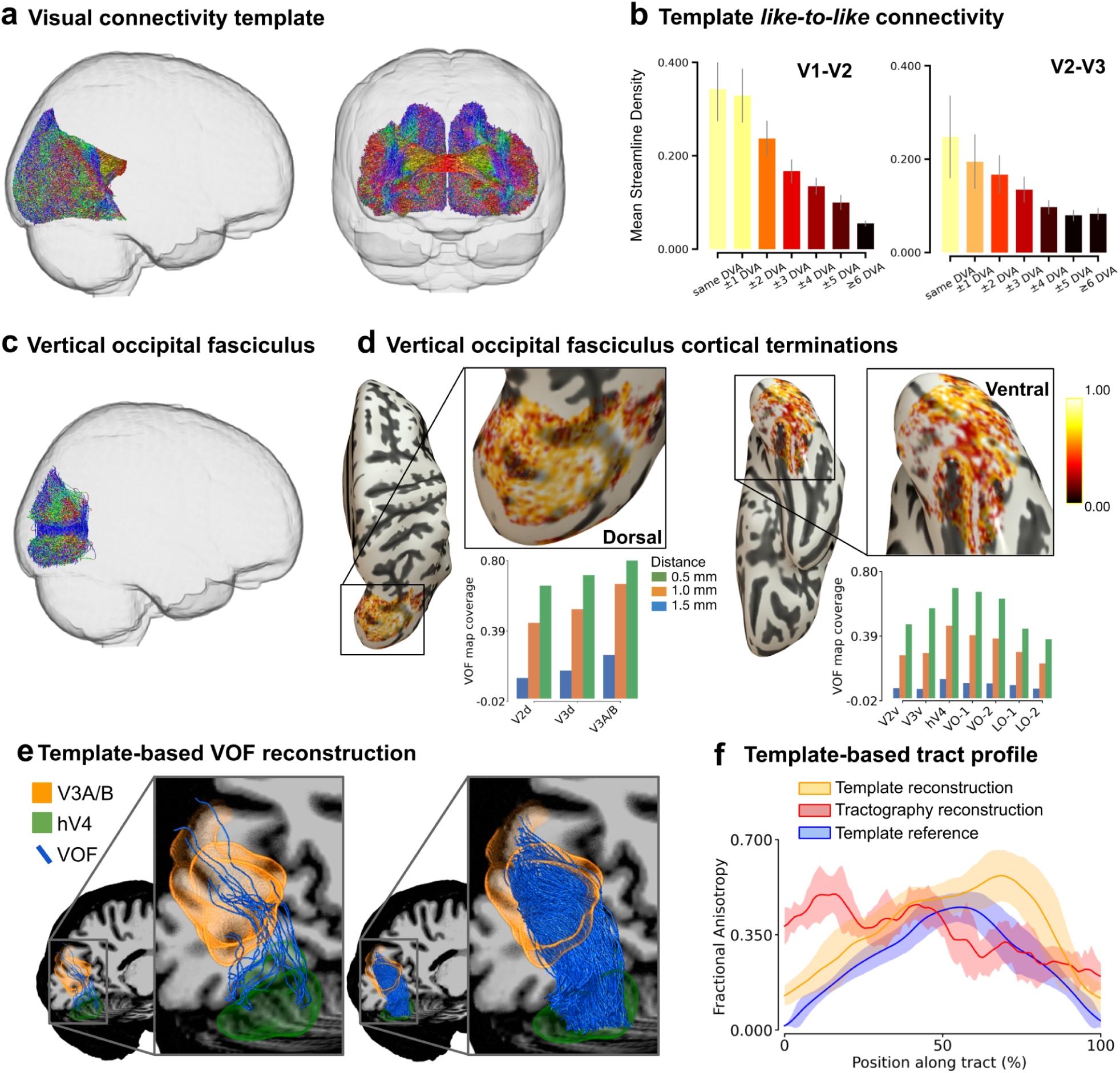
Population templates recover retinotopic connectivity beyond tractography limitations. **a.**, Population template of visual connectivity constructed from aggregated tractography across participants and represented in MNI space. Streamlines are colored according to retinotopic coordinates (eccentricity and polar angle), illustrating the geometric organization of visual pathways across multiple areas, including V1–V2, LO1–LO2, hV4, and V3A/B. **b.** Connectivity derived from the template preserves the like-to-like retinotopic organization. Eccentricity-by-eccentricity connectivity matrices for V1–V2 (*left*) and V2–V3 (*right*) show maximal connectivity between homologous visual field locations and decreasing connectivity with increasing eccentricity separation. **c-d.** The template reproduces known anatomical pathways. The vertical occipital fasciculus (VOF^30^) is segmented from the template (**c**), and its cortical termination patterns (**d**) match established anatomical organization. **e.** Template-based inference recovers connections that are not detected in individual tractography. Example of hV4–V3A/B connection: streamline reconstruction fails in a representative subject using standard tractography (*left*), but is recovered using the population template (*right*). **f.** Quantitative comparison of fractional anisotropy (FA) for the hV4–V3A/B pathway demonstrates consistent microstructural estimates following template-based reconstruction.

First, we tested whether *T_t_* preserves the retinotopic organization observed in individual-level analyses. Connectivity derived from *T_t_* reproduced the characteristic like-to-like pattern: streamline density was maximal between homologous eccentricity bins and decreased systematically with increasing eccentricity separation (**Fig. 4b**). This organization closely matched the population-level results (**Fig. 2**), indicating that *T_t_* retains the coordinate structure of retinotopic connectivity. To do so, streamline density (*S_d_*) was computed between all pairs of eccentricity bins in V1 and V2, forming eccentricity-by-eccentricity connectivity matrices (**Fig. 4b**, left). Results derived from *T_t_* reproduced the population-level pattern: connectivity was maximal along the main diagonal, corresponding to homologous eccentricity bins, and decreased systematically as eccentricity distance increased (**Fig. 4b**). We next assessed whether *T_t_*preserves known anatomical organization of visual pathways. Focusing on the vertical occipital fasciculus (VOF), a well-characterized tract with established cortical projections^30^ (**Fig. 4c**), we segmented the VOF from the template and quantified its cortical terminations (**Fig. 4c,d**). The resulting coverage map patterns reproduced the expected anatomical distribution, with strong projections to dorsal visual areas, including V3A/B, and comparatively weaker coverage in ventral early visual cortex, consistent with prior reports. These results indicate that the template retains known anatomical features of visual white-matter organization.

We then tested whether the template can recover connections that are not reliably reconstructed in individual tractography. In some participants, tractography failed to recover specific pathways, particularly for connections between hV4 and V3A/B (**Fig. 4e**, *left*), likely due to variability in cortical folding and local reconstruction limitations. Projecting the population template into individual anatomy restored these missing connections while preserving their retinotopic organization (**Fig. 4e**, *right*). Quantitative analysis of tract microstructure further supported this recovery. When tractography failed, fractional anisotropy (FA) estimates along the affected pathways deviated from the population distribution. In contrast, template-based reconstructions produced FA profiles are consistent with the population-level distribution while retaining individual participants variability (**Fig. 4f**), indicating that the recovered pathways reflect plausible anatomical structure rather than reconstruction artifacts. Together, these results demonstrate that population templates preserve key anatomical features of retinotopic connectivity and mitigate common tractography limitations, enabling robust inference of retinotopically specific pathways even when individual reconstructions are incomplete.

## Discussion

A central goal of neuroscience is to understand how anatomical connectivity links neural representations to behavior. Here we show that structural connectivity across the human visual cortex preserves retinotopic organization, such that cortical regions representing similar locations in visual space preferentially connect. By integrating diffusion tractography with maps of eccentricity and polar angle across large populations, the VISCONN framework reveals a retinotopic wiring principle linking cortical maps to the anatomical architecture of visual networks. These findings indicate that the geometry of visual space provides a scaffold for organizing long-range connectivity in the human brain.

Retinotopy has long been recognized as a defining organizational principle of the visual cortex. Classic electrophysiological studies demonstrated a systematic mapping of visual space across the striate cortex and neighboring extrastriate areas^74–76^. Modern neuroimaging has confirmed that these maps are highly reproducible across individuals and datasets, while simultaneously exhibiting systematic variation in cortical magnification and surface area^24,77,80,88^. However, the extent to which this spatial organization also constrains long-range anatomical connectivity had yet to be interrogated. The present results provide compelling evidence that it does.

Reconstructed structural pathways preferentially link cortical regions representing homologous locations in the visual field, suggesting that retinotopic coordinates provide a fundamental spatial scaffold for inter-areal connectivity. Together with previous findings linking retinotopic sampling to perceptual performance fields^15,24^, these results suggest that retinotopy organizes not only cortical representations but also the anatomical wiring that links visual areas into distributed functional networks. We note that connectivity was higher at the intercardinal locations in the lower visual field than in the upper visual field, although contrast sensitivity, spatial resolution and acuity are most pronounced at the vertical meridian and decrease gradually toward intercardinal locations, where they are no longer significant.^50,51,53,54,60,63,69^

These observations align with emerging principles from modern connectomics. Mesoscale reconstructions of mouse visual cortex demonstrate that projections between cortical areas preferentially connect neurons with similar tuning properties, revealing a “like-to-like” wiring rule governing inter-areal connectivity^12,13^. Complementary datasets from the MICrONS project similarly show that synaptic connectivity patterns across visual areas reflect functional similarity in neuronal responses^89^. Although diffusion MRI cannot resolve synaptic connections, the retinotopically aligned pathways observed here are consistent with the idea that macroscale anatomical connectivity reflects underlying functional similarity among neuronal populations. In this framework, the retinotopic connectivity patterns reconstructed may represent the mesoscale-level manifestation of circuit-level connectivity rules revealed in cellular connectomics.

Physiological studies in non-human primates provide converging support for this interpretation. Electrophysiological recordings demonstrate that receptive-field properties in macaque V2 emerge from spatially organized inputs originating from V1 neurons with aligned receptive-field locations and orientation preferences^90,91^. These findings indicate that structural connectivity is a substrate that shapes receptive-field computations performed by downstream cortical areas. The retinotopically aligned white-matter pathways identified here may therefore represent the macroscale anatomical infrastructure supporting such circuit-level transformations in the human brain.

Structural connectivity likely represents a key anatomical substrate for perception. We interrogated this link by investigating well-established asymmetries in visual perception. Behavioral studies have shown that perceptual performance varies systematically across polar angle and meridian locations of the visual field. In particular, contrast sensitivity^50–59^, spatial resolution and acuity^60–69^ are superior along the horizontal than the vertical meridian, and lower than the upper visual meridian. Neuroimaging studies have further demonstrated that these perceptual asymmetries correlate with variations in cortical magnification and retinotopic sampling in the early visual cortex in human adults^15,24,77,92^ and children^79^. Moreover, information accrual is faster along the horizontal than the vertical meridian, and along the lower than the upper vertical meridian^93^. For a review, see^19^. The connectivity patterns observed here raise the intriguing possibility that white-matter organization constitutes an additional anatomical component of these phenomena. If structural pathways preferentially connect cortical regions representing corresponding visual field locations, variations in pathway density or geometry could effectively amplify differences in cortical representation across the visual field, thereby constraining the information-processing capacity of specific meridians.

More broadly, these results align with recent work showing that brain networks are constrained by spatial geometry. Analyses of large-scale functional connectivity demonstrate that brain networks exhibit a topological structure reflecting spatial embedding within the folded cortex^94^. Rather than behaving as abstract graphs of discrete nodes, neural networks appear strongly constrained by anatomical geometry and minimization of wiring cost. The retinotopic connectivity patterns observed here may represent a domain-specific optimization of this broader principle, in which the geometry of visual space dictates the organization of structural connections between cortical areas.

Interpreting diffusion-based connectivity requires acknowledging the methodological limitations of tractography. Diffusion MRI measures water diffusion rather than axonal fibers directly; consequently, reconstructed streamlines represent statistical estimates of fiber trajectories rather than physical axons. This distinction is critical as tractography may generate both false-positive and false-negative connections. Large-scale validation studies demonstrate that even advanced tractography algorithms can produce anatomically implausible pathways under complex fiber configurations^86^. Comparisons between tractography and tracer-based connectivity further show that diffusion MRI may fail to recover a substantial fraction of known projections while generating spurious pathways^95^. Additional challenges arise from the organization of superficial white matter: dense sheets of short association fibers beneath the cortical surface can impede the ability of tractography algorithms to resolve long-range projections between cortical areas^96^. Diffusion-based connectivity estimates should therefore be interpreted as probabilistic models of anatomical connectivity rather than direct measurements of axonal projections^1,2^. The framework we present mitigates these limitations by constraining tractography with independently derived retinotopic constraints and by aggregating evidence across large populations into a template, thereby stabilizing inference about reproducible connectivity patterns.

The present work also illustrates the growing importance of population neuroimaging for studying human brain organization. Large datasets generated by initiatives such as the Human Connectome Project have enabled the construction of normative maps of structural and functional connectivity across the brain^42^. Complementary population studies, including PING and Cam-CAN, extend these approaches across development and aging^81,82^.

By aggregating tractography across a large cohort, the VISCONN template provides a high-fidelity population-derived representation of visual connectivity that captures invariant features of the visual connectome while maintaining a benchmark for quantifying individual variability.

Together, these findings demonstrate that the structural architecture of the human visual system reflects a fundamental organizational principle: cortical regions representing homologous locations in visual space preferentially connect with one another. By integrating retinotopic mapping with diffusion-based tractography across large populations, we reveal that the geometry of visual space dictates long-range anatomical connectivity in the human brain. This topographically specific wiring provides the anatomical substrate through which distributed visual areas coordinate concurrent processing of the same portion of visual space. Critically, we show that this wiring architecture is not uniform, but instead mirrors well-known perceptual asymmetries, suggesting that the density of structural connectivity constrains human visual performance. Ultimately, by bridging scales of analysis spanning cellular connectomics, systems neuroscience, and population neuroimaging, this study provides a structural scaffold supporting human visual function.

## Online Methods

### Data Sources

All data collections were approved by the respective local Institutional Review Boards (IRBs (see original papers for information)). Three publicly available neuroimaging datasets were utilized to examine the validity, reliability, and reproducibility of the results, as well as to characterize population-level distributions. Detailed information on image acquisition protocols, participant demographics, and preprocessing pipelines for each dataset can be found in the related publications^42,48,81,82^.

Data from the Human Connectome Project (HCP S1200)^42^ release were used. Structural MRI (sMRI): We used the minimally preprocessed structural T1-weighted (T1w) and T2-weighted (T2w) images from 1,066 participants in the HCP (S1200 release). Specifically, the 1.25 mm isotropic *“acpc_dc_restored”* T1w images acquired on a Siemens 3 T MRI scanner were included for all analyses involving HCP data. For our structural analyses, we leveraged the pre-processed *FreeSurfer*^34^ outputs provided by the HCP minimal preprocessing pipeline. Diffusion MRI (dMRI): We also incorporated the minimally preprocessed diffusion MRI data from the same 1,066 HCP participants (3 T Siemens scanner). All analyses utilized the full multi-shell diffusion acquisition data available in this release.

Data from the Cambridge Centre for Ageing and Neuroscience (Cam-CAN)^82^ study were used. Structural MRI (sMRI): We included the unprocessed structural T1w and T2w images (1 mm isotropic resolution) from 652 Cam-CAN participants. Diffusion MRI (dMRI): We also included the unprocessed diffusion-weighted images (2 mm isotropic resolution) from the same 652 participants in the Cam-CAN dataset.

Data from the Pediatric Imaging, Neurocognition, and Genetics (PING)^81^ study were used. Structural MRI (sMRI): We used the unprocessed structural T1w images (with voxel dimensions of 1.2 × 1.0 × 1.0 mm) and T2w images (1.0 mm isotropic) from 110 participants in the PING study. Diffusion MRI (dMRI): Similarly, we utilized the unprocessed diffusion MRI scans (2 mm isotropic resolution) from these 110 PING participants.

### Reproducibility and extensibility

All analyses were implemented as modular Apps in the *brainlife.io* platform^48^ and are publicly available with corresponding GitHub repositories. Each App is version-controlled and assigned a unique Digital Object Identifier (DOI), resolving to the specific instance used to generate the results reported here. This approach ensures full computational reproducibility by preserving the exact processing configuration and execution environment, while enabling extensibility through transparent, reusable components. Individual processing steps are referenced throughout the manuscript using their corresponding brainlife.io App identifiers (e.g., A0 or A116) and DOIs. A complete list of Apps, DOIs and source code GitHub repositories is provided in **Supplementary Table 6**.

### Data Preprocessing

#### Structural MRI pre-processing

HCP structural images were used as distributed (minimally preprocessed), with no additional structural preprocessing. For Cam-CAN, structural T1w and T2w images underwent bias-field correction and alignment to the anterior–posterior commissure (AC–PC) plane ([A273](https://doi.org/10.25663/brainlife.app.273) and [A350](https://brainlife.io/app/5ec6eda01f4464acbc9e54ec)). For PING AC–PC alignment was performed for both T1w and T2w images ([A99](https://doi.org/10.25663/bl.app.99) and [A116](https://doi.org/10.25663/brainlife.app.116)). Across datasets, participant-specific T1w volumes were segmented into tissue classes using the MRtrix3 function 5ttgen^36^([A239](https://doi.org/10.25663/brainlife.app.239)). The resulting gray–white matter interface mask served as the seed for white-matter tractography. T1w and T2w images were further processed with FreeSurfer recon-all^34^ for cortical and subcortical segmentation and surface reconstruction ([A0](https://brainlife.io/app/58c56d92e13a50849b258801)).

### Diffusion (dMRI) processing

#### Preprocessing and model fitting

For HCP S1200, the minimally preprocessed dMRI data were used without additional preprocessing. For Cam-CAN and PING, dMRI preprocessing followed the protocol in^97^ ([A68](https://brainlife.io/app/5a813e52dc4031003b8b36f9)). Briefly, data were denoised (dwidenoise) and corrected for Gibbs ringing (*mrdegibbs*)^36^, followed by correction of susceptibility-induced distortions, eddy currents, and head motion using FSL’s *topup* and *eddy*^35,98,99^. Eddy-current and motion correction was employed with outlier slice replacement (--repol)^100^. Gradient table consistency was verified and corrected with MRtrix3 *dwigradcheck* after *topup*/*eddy* ^101^. Volumes then underwent bias-field correction using ANTs N4^102^; residual background noise was additionally addressed with MRtrix3 *dwidenoise*^103^. Finally, diffusion volumes were registered to each participant’s T1w image using FSL *epi_reg*^104–106^. Brain masks for dMRI were generated with FSL *BET*^105,107^([A163](https://doi.org/10.25663/brainlife.app.163)). Following preprocessing, constrained spherical deconvolution (CSD) diffusion model^108^ was fit across spherical harmonic orders (Lmax = 2, 4, 6, and 8) using MRtrix3 ([A238](https://brainlife.io/app/5db224e98aeeee0f87f25c1f) and [A319](https://doi.org/10.25663/brainlife.app.319)).

#### Tractography

Whole-brain tractograms were generated using anatomically constrained probabilistic tractography (ACT) via the MRtrix3 toolkit^43^([A297](https://brainlife.io/app/5e88b8bf952fef39d57abcf2) and [A319](https://doi.org/10.25663/brainlife.app.319)). Fiber Orientation Distribution Functions (fODFs) were selected using a spherical harmonic representation, with the maximum harmonic degree (Lmax) chosen based on each dataset’s diffusion MRI acquisition. In all cases, we used 3,000,000 streamlines per subject, propagated with a 0.2 mm step size, and we enforced a maximum curvature of 35° to prevent anatomically implausible sharp turns. For HCP data, we used a higher spherical harmonic order (Lmax = 8) to estimate fODFs from the multi-shell diffusion data. Streamlines were filtered to retain lengths between 25 mm (minimum) and 250 mm (maximum), reflecting the adult brain dimensions in this cohort. Also, for PING data, we employed Lmax = 8 for fODF, but to accommodate smaller brain sizes in children and adolescents, we accepted streamline lengths from 20 mm to 220 mm. All other tracking parameters (3 million probabilistic streamlines, 0.2 mm step size) mirrored those of HCP. For Cam-CAN, we utilized Lmax = 6 for fODFs. The tractography produced streamlines of 25–250 mm in length (matching the HCP criteria), with the same 3,000,000 count and 0.2 mm step size as above. Across all datasets, the use of ACT ensured that tracking was guided by individual anatomical information (i.e., white and gray matter interfaces). These consistent tractography procedures provided comparable whole-brain streamline reconstructions while accounting for differences in data quality and subject anatomy.

### Population Receptive Field (pRF) Mapping and Retinotopic Parcellation

#### pRF Modeling and Visual Field Measures

Population receptive field (pRF) models were fitted to each participant’s cortical surface (derived from FreeSurfer^34^) using an automated procedure^40,41,109^([A559](https://brainlife.io/app/6115c829a5a04c2a80f9e7f7)). This method provided two primary retinotopic measures for each cortical vertex: *polar angle* (indicating the angular *quadrant* of the visual field to which that location responds; **Fig. 1a** *left*) and *eccentricity* (indicating the distance from the fovea, in degrees of visual angle, to which it responds; **Fig. 1a** *middle*). In addition to these continuous maps, an anatomical parcellation of visual areas was obtained for each participant based on the pRF results. Specifically, twelve visual cortical areas of interest were identified from the retinotopic mapping procedure, corresponding to early and mid-level visual areas: V1, V2, V3, V3A, V3B, hV4, LO1, LO2, TO1, TO2, VO1, VO2. Each of these areas was delineated by the pRF-based atlas, allowing region-specific analyses in subsequent steps.

#### Cortical Surface Map Generation for Polar Angle and Eccentricity

For both polar angle and eccentricity measures, the continuous maps were further processed into discrete “binned” maps on each participant’s cortical surface. This involved grouping the data into specified value ranges (bins, *b*; **Fig. 1a** *right*). The creation of these binned maps was accomplished using a combination of Connectome Workbench and FSL commands ([A784](https://doi.org/10.25663/brainlife.app.784)).

#### Iso-Eccentric Bin Definition

Iso-eccentric bins (*b_e_*) were defined by thresholding the eccentricity map into discrete intervals of visual field representation. At a resolution of 2° of visual angle (DVA), the visual field was partitioned into concentric eccentricity bands spanning 0–2° (foveal), 2–4°, 4–6°, and 6–8° (see **Fig. 1a**, *right*).

#### Polar Angle Meridian Definition

Meridians were defined within a selected polar angle range across the entire range of eccentricities, 0-90°. More specifically, 8 meridians were segmented for our analyses: i) Upper Vertical Meridian: 0° – 15°, ii) Lower Vertical Meridian: 165° – 180°, iii) Lower Vertical Diagonal: 120° – 150°, iv) Upper Vertical Diagonal: 30° – 60°, v) Horizontal Meridian: 75° – 105°. Each meridian identified a specific wedge on the retinotopic map (see **Fig. 3a**).

#### Tract Selection and Retinotopic Binning

To quantify retinotopic connections between visual areas, we restricted the streamlines to those connecting pairs of regions of interest (ROIs) defined within specific visual areas (e.g., V1 and V2). In this formulation, a retinotopic connection was defined as the set of streamlines linking one ROI in V1 to a corresponding ROI in V2, both constrained by eccentricity and polar angle bins (see **Fig. 1a**, *right*). For each polar angle meridian segment (e.g., Lower Vertical Meridian, 165°–180°), streamlines were subdivided according to the cortical eccentricity bins (e.g., 0–2°, 2–4°, 4–6°, 6–8°) to which their endpoints projected. To obtain retinotopically specific tracts, we first selected streamlines from the whole-brain tractogram (see **Fig. 1b**, *left*) that intersected cortical regions constrained jointly by the given eccentricity bin and polar angle sector (see **Fig. 1b**, *middle*). Streamlines whose endpoints fell within the same meridian and eccentricity bin were grouped to form iso-eccentric tract subsets, corresponding to fibers linking cortical representations of progressively more peripheral visual field locations (see **Fig. 1b**, *right*). For an example of the workflow, see **Supplementary Fig. 2**.

#### Retinotopic Tract Profiles

To quantify microstructural properties along retinotopic-defined connections, we computed Retinotopic Tract Profiles (*r_tp_*). Each *r_tp_*represents the variation of diffusion-derived metrics (e.g., Fractional Anisotropy) along streamlines connecting retinotopically defined cortical regions ([A911](https://brainlife.io/app/69ca22f834e13335187c6ae7)).

#### Tract Profile Computation

For each iso-eccentric tract group, diffusion metrics such as Fractional Anisotropy (FA) were sampled along the tract length using a streamline-based weighting approach, named *tract profile*^110^. FA reflects the degree of directional water diffusion within white matter and is commonly interpreted as a proxy for microstructural integrity, including factors such as axonal coherence and myelination. For each retinotopic tract, a centroid was computed from the streamline pathways^110^ and node-wise FA values were averaged across streamlines, weighted by their distance to the centroid^110^. Node-wise means were then computed relative to the tract centroid, which was estimated using a skeleton-based method^111^. In this approach, the centroid (the skeleton) was calculated by averaging the streamlines of the tracts belonging to the most dense portion of the tract itself, thresholding the streamlines passing through the 95% of the maximum density, ensuring that streamlines lying within the core fiber bundle (and not peripheral outliers) exert greater influence^111^. This weighting enhances the stability of the centroid estimate, especially in tracts with asymmetric or noisy spatial dispersion. During profile generation, each node value was weighted by the inverse of this Mahalanobis distance, providing smooth transitions across the tract and minimizing bias from sparsely represented or spatially deviating streamlines^110^.

#### Like-to-like retinotopic connectivity analysis

To quantify retinotopic specificity of structural connectivity between visual areas, we measured connectivity as a function of eccentricity between pairs of early visual cortical regions. Analyses focused on the area pairs V1–V2 and V2–V3, which share aligned retinotopic representations and well-characterized anatomical connectivity.

#### Definition of iso-eccentric cortical bins

For each visual area, the cortical surface was subdivided into iso-eccentric bins of 1° DVA, spanning the full eccentricity range analyzed in the study (0-20° DVA). Each bins, therefore, represented a cortical region encoding a narrow band of visual field eccentricity.

#### Streamline assignment to iso-eccentric bins

Whole-brain tractography streamlines were intersected with the gray–white matter interface of the retinotopically defined eccentricity bins. A streamline was considered to connect two eccentricity bins if its endpoints terminated within the corresponding gray–white interface voxels of those bins^36^.

#### Connectivity matrix construction

For each subject and each pair of visual areas, we computed an eccentricity-by-eccentricity connectivity matrix ([A910](https://brainlife.io/app/69ca2f0934e13335187cfbbd)). Each matrix element represented the structural connectivity between one eccentricity bin in the source area and one bin in the target area (see **Fig. 2b** *left*). Connectivity strength was quantified using a streamline density metric that normalized streamline counts by the size of the cortical regions involved. The calculation was performed as follows: For any two bins, the number of tractography streamlines connecting those two regions was counted. This raw streamline count was then normalized by the average volume of the two regions. Formally, for regions A and B, streamline density (SD) is defined as:

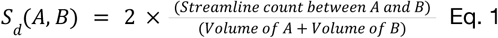

Using the mean of the two parcel volumes in the denominator yields a volume-normalized connectivity metric (streamlines per unit volume). This normalization controls both for differences in parcel size and reduces biases associated with larger cortical regions producing greater streamline counts, so correcting for the magnification effect of the visual maps on the cortex surface^75^. Connectivity matrices were computed independently for each participant and then averaged across subjects to obtain population-level estimates.

#### Diagonal-distance analysis

To quantify how connectivity varied as a function of eccentricity difference, elements of the eccentricity-by-eccentricity connectivity matrix were grouped according to their distance from the main diagonal. In this matrix, rows correspond to eccentricity bins in the source visual area and columns correspond to eccentricity bins in the target visual area. Each matrix element *a_ij_* therefore represents the streamline density between eccentricity bin *i* in the first area and eccentricity bin *j* in the second area. The *main diagonal* of the matrix contains elements for which the row and column indices are identical (e.g., *a_00_,a_11_,a_22_,a_33_*), corresponding to connections between homologous eccentricity bins across the two visual areas (i.e., identical eccentricity range that is the same degree of visual angle (DVA). Elements located away from the main diagonal represent connections between bins with increasing eccentricity difference. For example, the first off-diagonal consists of elements immediately adjacent to the main diagonal (e.g., *a_01_,a_12_,a_23_* above the diagonal and *a_10,_a_21_,a_32_* below the diagonal), corresponding to connections between eccentricity bins differing by one bin (+/- 1 DVA). Similarly, the second off-diagonal contains elements such as *a_02_, a_13_, a_24_,* and *a_20_, a_31_, a_42_*, representing connections between bins differing by two eccentricity steps (+/- 2 DVA). To obtain a summary measure of connectivity as a function of eccentricity difference, elements belonging to the same diagonal offset were grouped. Specifically, elements from the upper and lower off-diagonals at the same distance from the main diagonal (e.g., *a_01_, a_12_, a_23_*, and *a_10_, a_21_, a_32_*) were averaged to produce a single estimate of connectivity for that eccentricity difference. This procedure produced a one-dimensional profile describing how streamline density varied as a function of eccentricity difference between the connected visual regions (see **Fig. 2b** *right*). The diagonal-distance analysis was applied to the aggregated matrices to quantify how connectivity strength varied as a function of eccentricity difference between connected visual areas.

### Visual Connectivity Asymmetries analysis

#### Meridian-based ROI definition and tract selection

To investigate visual field asymmetries in structural connectivity, analyses were restricted to polar angle–defined regions of interest (ROIs), independent of eccentricity. Early visual areas (e.g., V1, V2) were subdivided into wedge-shaped ROIs corresponding to the major visual field meridians: horizontal meridian (HM), upper vertical meridian (UVM), and lower vertical meridian (LVM) (see **Fig. 3a** *left*). Each ROI encompassed all eccentricities within a restricted polar angle range centered on the corresponding meridian. Whole-brain tractograms were then filtered to isolate streamlines whose endpoints terminated within matched meridian ROIs across visual areas. For example, V1–V2 horizontal meridian connectivity was defined as the set of streamlines with one endpoint in the HM ROI of V1 and the other in the HM ROI of V2. The same procedure was applied to the UVM and LVM ROIs, yielding homologous meridian-to-meridian connection sets. This approach enabled direct comparison of structural connectivity strength across different regions of the visual field. Two categories of structural connections were defined: (1) short-range (local) connections: both endpoints of a streamline terminate within visual area ROIs belonging to the same iso-eccentric, polar angle–defined meridian “wedge” in the occipital lobe (see **Fig. 3a** *right*)([A784](https://brainlife.io/app/64842d58b10869ed4e8bbd16)); (2) long-range (global) connections: only one endpoint of a streamline lies within an occipital iso-eccentric meridian visual area, while the other endpoint is located outside the occipital parcellation. Thus, the streamline connects a retinotopically defined occipital region to a different brain area (e.g., a non-visual cortical lobe; see **Fig. 3c**)([A785](https://brainlife.io/app/648d4de5b10869ed4e9ada5b)).

#### Streamline Density Computation

For each pair of ROIs, structural connectivity strength was quantified using a *S_d_* metric (as defined in *Eq. 1*). For long-range (global) connections, streamline density was computed on a per-region basis. For each occipital visual area parcel, we counted the total number of long-range streamlines terminating in that region (i.e., streamlines that originate outside the occipital lobe and end within the given visual region). This count was then divided by the volume of that visual region to yield a normalized measure of incoming long-range connectivity density (streamlines per unit volume). This metric reflects how densely a given visual area is connected to distant brain regions, adjusted for the size of the region.

#### Quantification of visual field asymmetries

Meridian-specific streamline densities were compared to characterize asymmetries in structural connectivity across the visual field. In particular, we contrasted connectivity between horizontal and vertical meridians (HM vs. VM, where VM is defined as the union of UVM and LVM) to assess the horizontal–vertical asymmetry (HVA), and between lower and upper vertical meridians (LVM vs. UVM) to assess the vertical meridian asymmetry (VMA). These comparisons directly test whether structural connectivity between visual areas varies systematically as a function of polar angle wedges (see **Fig. 3a** *right*).

#### Structural Asymmetry Metrics

Two structural asymmetry indices were derived from the meridian-specific streamline densities. The horizontal–vertical asymmetry (HVA) index was defined as

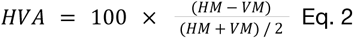

and the vertical meridian asymmetry (VMA) index was defined as

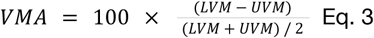

These normalized difference measures quantify the relative imbalance in streamline density between horizontal and vertical meridians (HVA) and between lower and upper vertical meridians (VMA), while controlling for the overall magnitude of the measurements.

#### Structural Asymmetry Analysis Across the Lifespan

To characterize the developmental and aging-related trajectories of the visual asymmetries, we performed a cross-sectional lifespan analysis of structural connectivity asymmetries across visual meridians. The aim is to quantify how structural visual field asymmetries (the HVA and VMA asymmetries) change from early childhood through late adulthood. Measurements from all cohorts were concatenated while preserving dataset identity. For each subject and meridian, streamline density (*S_d_*) was calculated as the number of streamlines connecting the meridian-specific region in one visual area to the corresponding region in the other, normalized by the mean volume of the connected areas.

#### Lifespan Modeling of Asymmetry Indices

Age-related changes in structural asymmetry were characterized using polynomial regression models relating age to each asymmetry metric. For each dataset independently, second-order polynomial models of the form

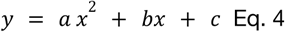

were estimated using least-squares regression, where (*y*) represents either HVA or VMA and (*x*) denotes participant age in years. Quadratic models were selected to capture potential nonlinear lifespan trajectories, including developmental increases and age-related decline. Models were estimated separately for each cohort (PING, HCP, and Cam-CAN) to maintain within-dataset consistency and to account for differences in acquisition protocols and preprocessing procedures. In addition, a global model was estimated using pooled subject data across cohorts to provide an overall lifespan trajectory. Model performance was quantified using the coefficient of determination (R^2^) and the Akaike Information Criterion (AIC).

#### Gyral Bias Analysis

To quantify the relationship between cortical folding and the distribution of visual field meridian representations, we performed a gyral bias analysis ([A914](https://brainlife.io/app/69cb8d9c34e133351885ac23)). This analysis investigated whether streamlines associated with specific meridians preferentially terminate in gyral versus sulcal regions of the cortex (i.e., the gyral bias), thereby linking tractography-derived connectivity measures to cortical geometry^112,113^. See also **Supplementary Results 6**.

#### Curvature Computation

Vertex-wise cortical curvature estimates were derived from FreeSurfer outputs (*curv* files), where negative values indicate gyral crowns and positive values indicate sulcal fundi. For each iso-eccentric meridian-defined parcel, curvature values were averaged across all surface vertices included in the parcel mask. This yielded a mean curvature measure per subject, per meridian, per eccentricity bin. The sign convention (gyri < 0, sulci > 0) followed the FreeSurfer standard.

#### Surface-to-Voxel Mapping of Curvature

To relate streamline terminations (voxel-level) to cortical folding (surface-level), vertex-wise curvature values from FreeSurfer were projected to the gray–white matter interface voxels. This mapping was performed by interpolating each vertex’s curvature value to its corresponding voxel in the subject’s structural space using FreeSurfer’s surface-to-volume tools (*mri_surf2vol*). As a result, each voxel at the cortical surface was assigned a curvature value consistent with the subject’s surface reconstruction, enabling voxel-wise correspondence between tractography endpoints and gyral/sulcal geometry.

#### Streamline Density Mapping

Streamline endpoints were intersected with the iso-eccentric meridian parcellations (see **Streamline Density Computation**). For each parcel, we computed the streamline density as the number of streamlines terminating within the parcel divided by the number of gray–white matter interface voxels in that parcel. To account for inter-subject differences in overall tractogram size, we further normalized these densities by the total number of streamlines generated for each subject.

#### Aggregated Analysis

For each subject and meridian, we aggregated voxel-level measures into parcel-level means, resulting in per-subject estimates of: (a) mean curvature (gyrus vs. sulcus tendency), (b) streamline density (corrected for parcel volume). Subject-level measures were averaged across participants to derive group-level centroids, while the standard error of the mean (SEM) was used to characterize inter-subject variability.

#### Density Estimation and Visualization

To characterize the joint distribution of streamline density and cortical curvature, we applied bivariate kernel density estimation (KDE) to subject-level data pooled across all participants (see **Supplementary Fig. 8**). Density estimates were computed separately for each meridian using a Gaussian kernel, and visualized as filled contour maps (five levels) to represent regions of increasing probability density. A lower density threshold (0.05) was applied to suppress low-probability regions and improve interpretability of the distributions. These KDE maps provide a smoothed representation of the curvature–connectivity relationship across subjects, rather than reflecting individual-level variability.

#### Scalable Generalized Streamline-based Algorithm for Tractogram Template Generation

A central contribution of this work is the construction of a population-level retinotopic template of visual white-matter connectivity derived from tractography in 1,049 individuals. This template provides a standardized reference that captures the typical spatial organization of visual pathways across the human brain (**Fig. 4a**). To generate the template, we aggregated occipital streamlines from participants in the Human Connectome Project, yielding an ensemble tractogram comprising more than 160 million streamlines.

Constructing a population-level tractography template requires addressing two fundamental challenges: the extreme scale of ensemble tractograms and the absence of a well-defined population average for streamlines. Even when restricted to the occipital lobe, each subject contributes on the order of ∼1.5 × 10⁵ streamlines, yielding an aggregate ensemble exceeding 10⁸ fibers across the population. Direct processing at this scale is computationally intractable on conventional hardware, necessitating a hierarchical, streaming-based strategy that preserves anatomical structure while ensuring bounded memory usage. Moreover, unlike voxel-wise measurements defined on a common grid, tractography streamlines are extended three-dimensional trajectories, precluding simple averaging across subjects and requiring geometric clustering and consensus estimation to derive representative pathways.

In a nutshell, to enable scalable processing while preserving anatomical structure, we developed a hierarchical framework that compresses this ensemble into a compact set of representative pathways. Because each participant contributed more than 150,000 occipital streamlines, the resulting collection contained over 160 million streamlines, posing substantial computational challenges. To reduce dimensionality while preserving anatomical structure, we designed a hierarchical clustering framework. First, we computed the union of all occipital streamlines across participants, generating an ensemble tractogram (*Tₑ*). Streamlines from Tₑ were randomly partitioned into 20 non-overlapping folds of approximately 8 million streamlines each. Within each fold, 80,000 clusters were computed using a dissimilarity embedding approach^114,115^, reducing dimensionality while preserving indices linking clusters to the original streamlines. From each cluster we extracted a medoid streamline, yielding approximately 1.6 million representative streamlines across folds. These medoids were then projected back into the full *Tₑ* space, where a searchlight algorithm computed a density-weighted average of the local streamline neighborhood surrounding each medoid^111^. The resulting tractogram was subsequently refined through anatomical masking and density-based pruning to remove weakly supported streamlines trajectories. This procedure produced a compact population template, a tractogram in MNI space containing approximately 1.5 million anatomically supported streamlines (**Fig. 4a**). Below we provide a more detailed procedure.

#### Ensemble construction and preprocessing

We constructed a population-level occipital tractogram using diffusion MRI data from 1,061 participants in the Human Connectome Project. Individual tractograms were spatially normalized to MNI space using nonlinear transformation derived from structural MRI registration^38,111,116^. Streamlines were restricted to those terminating within retinotopically defined visual cortical regions^40^, and implausible geometries and short fibers were removed^37,117^ ([A912](https://brainlife.io/app/69ca07bd34e13335187bffbc)). Quality control based on streamline counts and z-score outlier detection excluded 12 subjects, yielding 1,049 tractograms that were concatenated into a unified ensemble (∼163 million streamlines).

#### Streaming and fold-based data management

To ensure computational scalability, the ensemble was processed using sequential streaming of indexed chunks (20 million streamlines per chunk), limiting memory usage while preserving full dataset coverage. Streamlines were randomly redistributed into 20 balanced folds, each representing a statistically homogeneous subsample of the global distribution. This strategy ensures reproducibility, prevents ordering biases, and enables independent fold-wise processing.

#### Hierarchical clustering and medoid representation

Within each fold, streamlines were embedded in a dissimilarity space using orientation-invariant distances to prototype fibers^114,115^. Clustering was performed using MiniBatch K-Means^118^, partitioning each fold into 80,000 clusters ([A916](https://brainlife.io/app/69ca07bd34e13335187bffbc)). Rather than using centroids, which do not correspond to anatomically valid fibers, we extracted medoids defined as the streamlines minimizing within-cluster dissimilarity. Aggregating medoids across folds yielded a compact representation of approximately 1.6 million streamlines that approximates the full ensemble while preserving anatomical variability.

#### Searchlight refinement and skeleton estimation

The medoid representation was refined using a spatial searchlight procedure to estimate population-consistent anatomical pathways. White matter space was partitioned into spatial tiles constrained by tractography-derived density masks. For each medoid, neighboring streamlines within a fixed-radius sphere were identified, and representative “skeleton” streamlines were computed using density-weighted averaging^111^ ([A915](https://doi.org/10.25663/brainlife.app.915)). Streamlines were resampled to 100 points to ensure correspondence, smoothed using spline interpolation, and constrained to high-density regions (≥95% of local density) with minimum length thresholds (≥25% of maximum streamline length). This procedure suppresses noise and emphasizes the core geometry of white-matter pathways.

#### Template assembly and anatomical constraints

Refined skeleton streamlines were concatenated to form the population template. A final post-processing stage enforced anatomical validity by removing streamlines extending beyond the brain mask and pruning low-density trajectories. The tractogram was further partitioned into left-hemisphere, right-hemisphere, and commissural components using anatomical masks, enabling hemisphere-specific analyses. The final template comprises 1,539,761 streamlines and preserves the geometric organization of visual white-matter connectivity while remaining computationally efficient.

#### Template-driven retinotopic tract segmentation and subject-space projection

To enable subject-level inference of retinotopically specific pathways from the population template, we implemented a framework that consists of two sequential components: segmentation of retinotopically defined tract segments in template space, followed by their projection into subject-specific anatomical space (A913).

#### Visual tract extraction

To enable anatomically specific interrogation of white-matter connectivity, the framework incorporates a template-driven mechanism for extracting retinotopic tract connections. Using the Benson retinotopic cortical atlas^40,41^ in MNI space, eccentricity, polar angle, and visual-area labels are thresholded to generate regions of interest corresponding to user-defined retinotopic bins. Streamlines from the population occipital template are then filtered according to ordered endpoint inclusion within these ROIs, optionally constrained to individual hemispheres. This procedure yields tract segments that preserve true end-to-end anatomical pathways between retinotopic subregions rather than partial streamline fragments, ensuring biological interpretability of connectivity estimates. Because the segmentation is performed directly in MNI space, it remains independent of subject-specific noise and artifacts while retaining compatibility with downstream subject-level projection.

#### Registration-agnostic transformation to subject anatomy

Our procedure applies spatial transformations (linear or non-linear) to project retinotopic tractography streamlines from the template to individuals’ brain space. More specifically, we computed nonlinear registration between individual participants’ T1-w images and the MNI152 T1-w template^38^ ([A861](https://doi.org/10.25663/brainlife.app.861)). The resulting affine matrix and warp field are subsequently applied to the retinotopic tracts^37^ ([A902](https://brainlife.io/app/69aaf3006bb1f71b25ac8e59)), preserving streamline geometry under nonlinear deformation^111,116^.

#### Vertical Occipital Fasciculus (VOF) Segmentation and Coverage Quantification VOF candidate extraction and geometric refinement

The vertical occipital fasciculus (VOF) was isolated from whole-brain tractography using a waypoint-based segmentation procedure closely following the strategy described by Takemura et al.^30^ ([A917](https://brainlife.io/app/69cc4e4934e13335188d9217)), see **Fig. 4c,d**. Candidate streamlines were required to traverse two anatomically defined occipital waypoint planes aligned with the superior–inferior axis of posterior white matter, while hemisphere-specific exclusion masks prevented contamination from contralateral fibers. After waypoint filtering, candidate streamlines underwent geometric refinement to approximate the canonical vertical trajectory of the VOF. Streamlines were retained only if they satisfied constraints on minimum length, trajectory straightness, and orientation relative to the inferior–superior axis, consistent with the morphological criteria used in prior VOF characterizations. When specified, these constraints were evaluated within a bounded z-segment corresponding to the core vertical portion of the tract, thereby excluding oblique fibers intersecting the waypoint planes but not belonging to the principal vertical bundle. This procedure yielded a geometry-filtered VOF core for each hemisphere.

#### Retinotopic termination mapping and coverage estimation

To quantify the spatial relationship between the VOF and retinotopically defined cortical regions, streamline terminations from the filtered VOF bundle were evaluated relative to pRF-derived visual-area parcels. Following previous methods^30^, we computed the map coverage of an area given a tract as the proportion of cortical voxels within a given visual area that lie within a specified Euclidean distance of any VOF termination. Map coverage was computed across multiple distance thresholds (0.5, 1, and 1.5 mm), producing distance-dependent coverage profiles for each ROI (**Fig. 4 d**). In addition, a continuous coverage score map was generated by averaging binary coverage across thresholds, yielding voxelwise values in the interval [0,1][0,1][0,1] that summarize the spatial support of VOF terminations within retinotopic cortex (**Fig. 4 d**). Continuous maps were projected to the cortical surface for visualization and group-level comparison (**Fig. 4 d**). ROI-level metrics were averaged across hemispheres, yielding symmetric population estimates of VOF coverage and termination density suitable for group-level statistical analysis.

## DATA AVAILABILITY

All data generated and analysed during this study are publicly available through the brainlife.io platform as curated “Publications”. Derived datasets produced as part of this work are openly accessible at https://doi.org/10.25663/brainlife.pub.67.

User data agreements are required for some projects, like data from the Human Connectome Project (HCP), Cambridge Centre for Ageing and Neuroscience (Cam-CAN), and Pediatric Imaging, Neurocognition, and Genetics (PING) datasets.

## CODE AVAILABILITY

A complete record of all computational workflows used in this study, including brainlife.io Apps, their associated Digital Object Identifiers (DOIs), and corresponding GitHub repositories, is provided in Supplementary Table 1. The DOIs resolve to versioned instances of each App as executed on brainlife.io, ensuring full transparency and reproducibility of the analyses.

## ETHICS DECLARATIONS

The authors declare no competing financial interests.

## Acknowledgements

This research was supported by the following grants: Wellcome Trust (grant no. 226486/Z/22/Z, Principal Investigator F. Pestilli); NINDS UM1NS132207, BRAIN CONNECTS: Center for Mesoscale Connectomics (Principal Investigator K. Ugurbil); and NINDS U24NS140384, BRAIN CONNECTS: The Axonal Projectome EXchange (APEX) (Principal Investigator F. Pestilli). We thank Amazon Web Services Open Data Sponsorship Program for supporting data storage for brainlife.io

## SUPPLEMENTARY INFORMATION

### Supplementary Results 1. VISCONN pipeline

We developed VISCONN (VISion CONNectivity), an automated framework that integrates diffusion MRI tractography with retinotopic maps to estimate structural connectivity between cortical locations defined in visual field coordinates (see **Supplementary Fig. 1**). VISCONN combines anatomically predicted retinotopic maps with whole-brain tractography to reconstruct connections between cortical regions representing corresponding positions in visual space.

The pipeline operates on structural (T1-weighted) and diffusion-weighted MRI data acquired from each participant. Structural images undergo standard preprocessing such as AC (anterior commissure) - PC (posterior commissure) alignment and bias field correction^1^ and cortical surface reconstruction^2^. Retinotopic maps were automatically mapped as continuous estimates of polar angle and eccentricity derived from population receptive field (pRF) modeling using a Bayesian approach (*neuropythy*^3–5^). These maps were used to segment twelve visual areas (V1, V2, V3, V3A, V3B, hV4, LO1, LO2, TO1, TO2, VO1, VO2), each of which was further subdivided into discrete regions according to visual field coordinates. Specifically, cortical vertices were grouped into eccentricity bins and polar-angle sectors, enabling a spatially resolved representation of the visual field across the cortex. Diffusion-weighted MRI data were first preprocessed using standard pipelines implemented in FSL and MRtrix3 to correct for noise, motion, and imaging distortions^6–8^.

Fiber orientation distributions were then estimated using constrained spherical deconvolution to model the underlying white-matter fiber architecture^8,9^. Whole-brain anatomically constrained probabilistic tractography was subsequently performed to reconstruct white-matter pathways^10^, with streamlines generated at the individual subject level. Retinotopic parcellations were combined with tractography by assigning streamline endpoints to corresponding retinotopic parcels across visual areas, enabling identification of connections as a function of homotopic visual field position (see **Supplementary Fig. 2**). In this way, VISCONN enables estimation of connectivity between cortical locations representing the position in visual space, rather than between anatomically defined regions alone.

The output of the VISCONN pipeline consists of two complementary representations of structural connectivity. First, connectivity can be summarized as retinotopic connectivity matrices, which quantify streamline density between pairs of retinotopic regions across visual areas. These matrices are indexed by eccentricity and polar angle, enabling direct assessment of spatially specific connectivity patterns. Second, the microstructural properties of reconstructed pathways can be characterized using retinotopic tract profiles, which describe the variation of diffusion-derived metrics (for example, fractional anisotropy) along pathways linking specific visual field locations^11^.

VISCONN is implemented as a modular and fully automated workflow and is distributed as an open-source, containerized service on *brainlife.io* ^12^, enabling reproducible and scalable analyses across large neuroimaging datasets.

**Supplementary Figure 1.**
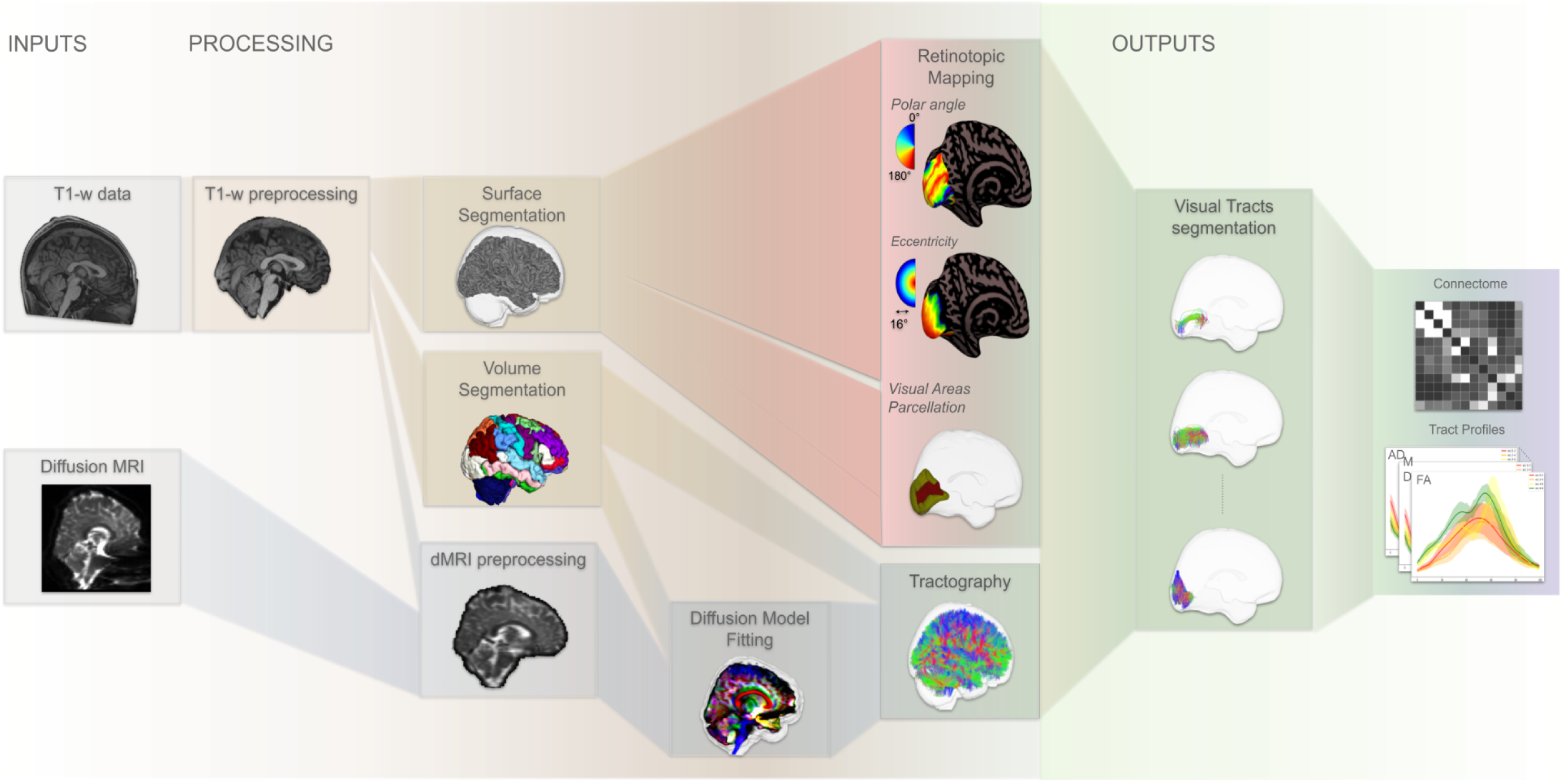
VISCONN pipeline for mapping retinotopic structural connectivity in the human visual system. Schematic overview of the end-to-end processing workflow implemented in VISCONN. Left (Inputs): Structural T1 -weighted MRI and diffusion-weighted MRI are acquired from each participant T1-weighted data undergo preprocessing (ANTs) and cortical surface reconstruction (Freesurfer), while diffusion data are preprocessed (FSL/MRtnx3) and used for whole brain tractography. Middle (Processing): Cortical surfaces are segmented into visual areas and further parcellated according to retinotopic coordinates (polar angle and eccentricity; Neuropythy). In parallel, diffusion models are fitted and ensemble tractography is performed (MRtri3/Dipy). Retinotopic parcellations are then combined with tractography to identify and quantify white matter streamlines connecting visual areas as a function of homotopic visual field position. Right (Outputs): The pipeline yields retinotopy-resoIved visual connectivity matrices and tract-specific microstructural profiles, enabling quantitative analyses of connectivity strength and asymmetries across eccentricity and polar angle. The workflow is fully automated, modular and containerized, and is deployed as an open-source service on brainlife.io4 to ensure reproducibility and scalability.

**Supplementary Figure 2.**
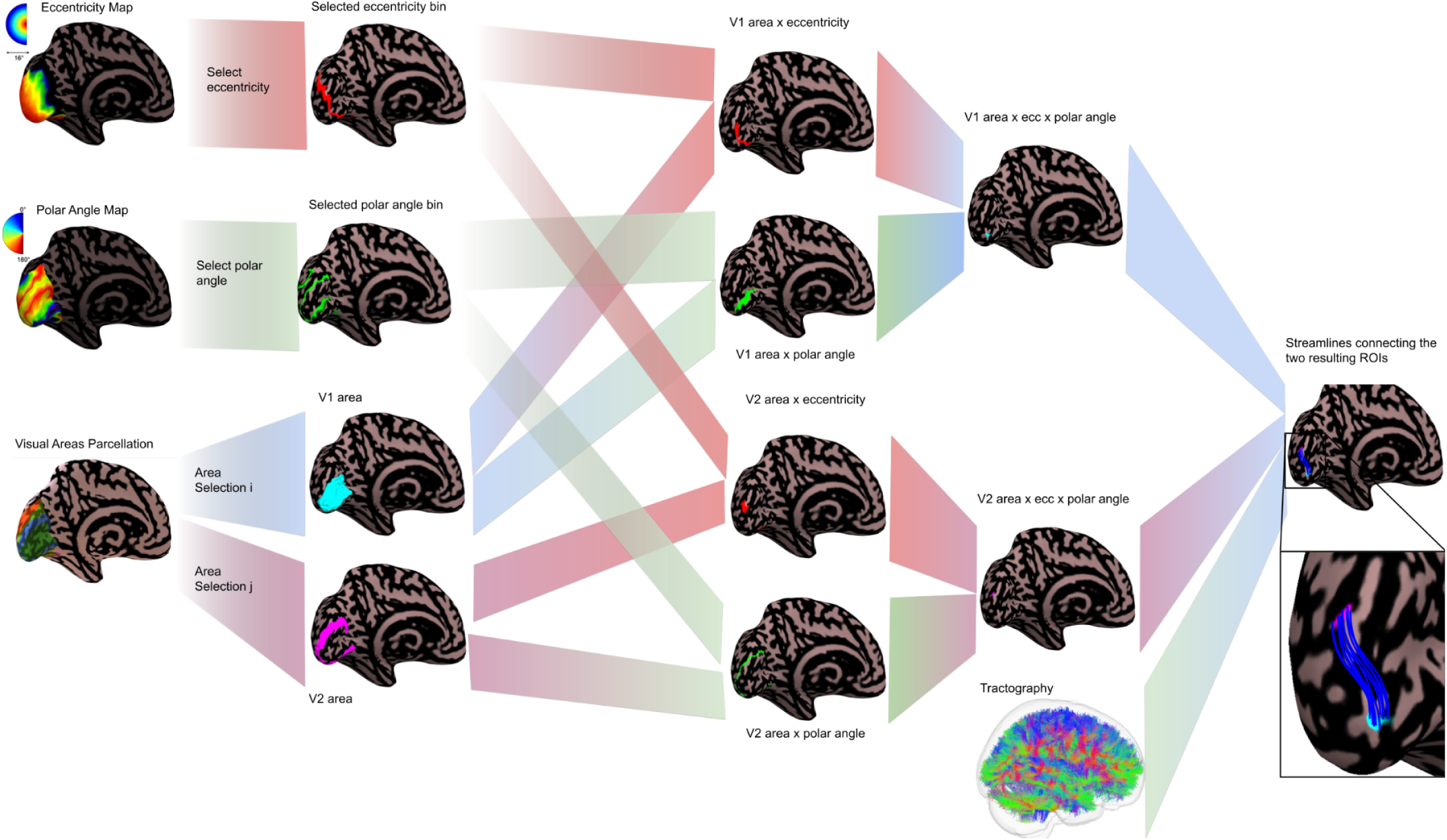
Workflow of VISCONN for selecting retinotopically constrained connections. VISCONN integrates retinotopic maps (eccentricity and polar angle) with visual area parcellations to isolate connections between specific cortical regions. First, the user selects an eccentricity range (eg, 4-6° dva) and a polar angle sector (eg, 75°-105°, corresponding to the horizontal meridian for a 15° wedge). In parallel, regions of interest (ROIs) are defined based on visual area parcellation (e.g., V1 and V2).These selections are combined to generate retinotopically constrained sub-ROIs (e.g., V1 x eccentricity × polar angle and V2 × eccentricity × polar angle). Finally, these sub-ROIs are intersected with whole-brain tractography to extract streamlines specifically connecting the selected visual areas within the defined retinotopic bins.The diagram illustrates each step of this selection and combination process, from input maps and ROI definition to the extraction of retinotopically specific connections.

### Supplementary Results 2. Retinotopic connectivity across visual areas as a function of eccentricity

To characterize how structural connectivity varies across the visual hierarchy as a function of eccentricity, we computed connectivity matrices across 12 visual areas (e.g., V1, V2, V3, and higher-order regions) separately for each eccentricity bin. For each subject and each eccentricity bin (0–2°, 2–4°, 4–6°, 6–8° DVA), we extracted the subset of streamlines whose endpoints fell within the corresponding eccentricity-defined ROIs. Using these subsets, we computed 12×12 connectivity matrices in which each element represents the streamline density (*Sd*) between a pair of visual areas constrained to that specific eccentricity bin. **Supplementary Fig. 3** illustrates these matrices across eccentricity bins: panel (a) shows the 0–2° matrix, (b) the 2–4° matrix, (c) the 4–6° matrix, and (d) the 6–8° matrix. Across bins, the matrices reveal systematic variations in connectivity strength across the visual hierarchy.

**Supplementary Figure 3.**
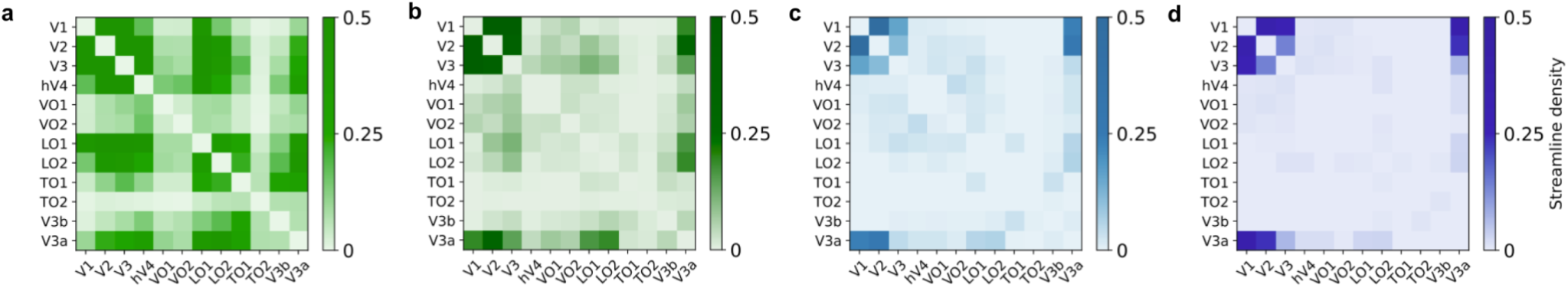
Eccentricity-resolved connectivity across visual areas. Connectivity matrices (12×12) showing streamline density (Sd) between visual areas for each eccentricity bin. (a) 0–2°, (b) 2–4°, (c) 4–6°, and (d) 6–8° (DVA). Each matrix element represents the connectivity strength between a pair of visual areas, computed using streamlines whose endpoints fall within the corresponding eccentricity range. This analysis characterizes how structural connectivity is distributed across the visual hierarchy as a function of eccentricity.

Connectivity patterns within foveal representations (0–2°; **Supplementary Fig. 3a**) show relatively strong and structured interactions among early visual areas, consistent with the high cortical magnification and dense connectivity associated with central vision. As eccentricity increases (**Supplementary Figb–d**), overall connectivity strength tends to decrease, and the distribution of connections across areas becomes more diffuse. This pattern is consistent with the reduced cortical representation and increased receptive field sizes associated with more peripheral visual field locations. Moreover, connectivity between early visual areas (e.g., V1–V2–V3) remains prominent across all eccentricity bins, whereas connections involving higher-order visual areas show greater variability across eccentricities. This suggests that while early-stage processing maintains relatively stable inter-area coupling across the visual field, higher-level regions may differentially integrate information depending on eccentricity.

### Supplementary Results 3. Control for retinotopic regions distance

It is established that for most tractography algorithms, the ability to track between two regions depends on the distance between the two regions^13,14^. To assess the influence of tract length on connectivity estimates, we recomputed retinotopic connectivity matrices after normalizing streamline density (S̅ _d_) by the mean length of streamlines connecting each pair of eccentricity-defined regions of interest. This normalization down-weights long-range connections relative to short-range connections, thereby controlling for distance-dependent biases inherent to tractography.

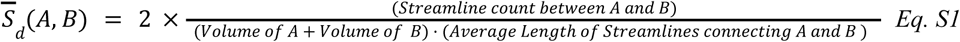

**Supplementary Figure 4.**
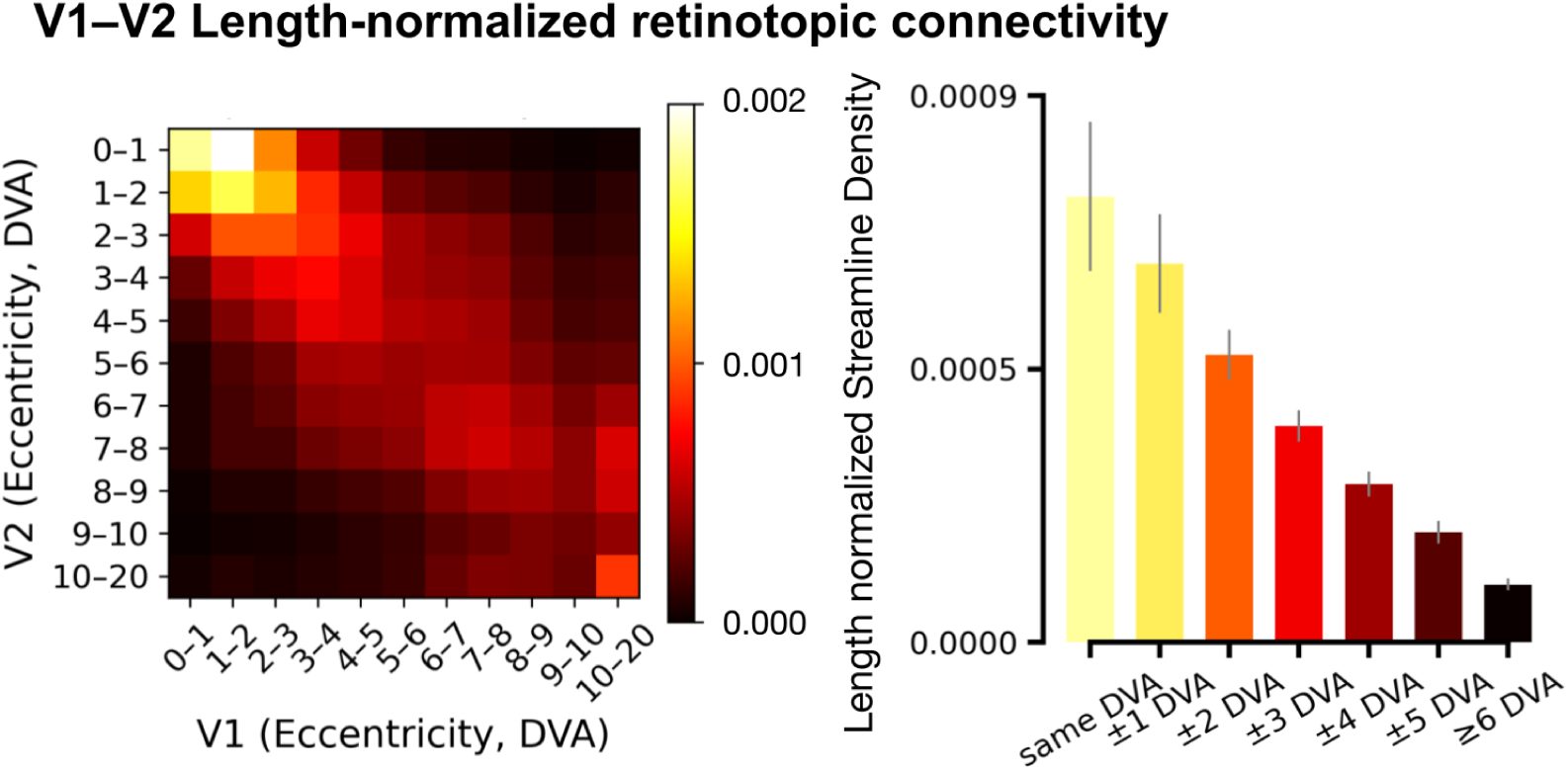
Length-normalized retinotopic connectivity. *Left.* Group-averaged V1–V2 retinotopic connectivity matrix after normalization of streamline density by both region surface area and mean streamline length for each eccentricity-bin pair. Connectivity remains maximal along the main diagonal, indicating preferential coupling between iso-retinotopic locations, consistent with the like-to-like organization observed in the non-normalized data (Fig. 2). *Right.* Mean *S* as a function of eccentricity distance, demonstrating maximal connectivity between matching eccentricity bins (same degrees of visual angle, DVA) and a monotonic decline with increasing separation. Length normalization reduces overall connectivity magnitude and attenuates long-range (off-diagonal) contributions, but preserves the topographic structure of the matrix. This indicates that the observed retinotopic organization is not driven by distance-dependent tractography biases.

Comparison of like-to-like connectivity patterns obtained without (**Fig. 2**) and with length normalization (**Supplementary Fig. 4**) revealed a substantial reduction in absolute connectivity values, but preservation of the underlying matrix structure (Pearson *r* = 0.994; Spearman ρ = 0.990). Row-wise analyses further showed that eccentricity-dependent connectivity profiles were largely unchanged (mean Pearson *r* ≈ 0.997), indicating that normalization induces a global rescaling without altering the relative organization of connections.

These results demonstrate that the like-to-like retinotopic organization is not explained by streamline length biases but reflects a robust topographic property of structural connectivity.

### Supplementary Results 4. Retinotopic connectivity asymmetry across early visual areas

To characterize how structural connectivity varies as a function of visual field location, we quantified streamline density between retinotopically defined regions across early visual areas and summarized these estimates as polar-angle–resolved connectivity profiles. Connectivity was evaluated across the four principal visual meridians: horizontal meridian (HM), vertical meridian (VM), lower vertical meridian (LVM), and upper vertical meridian (UVM), as defined from retinotopic maps derived using population receptive field (pRF) modeling^3–5^.

Connectivity profiles were visualized using radial (radar) plots, in which each axis represents a visual meridian, and values correspond to mean streamline density across participants (see **Supplementary Fig. 5**). Streamline density was computed as a volume-normalized measure of connection strength between retinotopic regions, enabling comparison across cortical locations with different surface area representations.

**Supplementary Figure 5.**
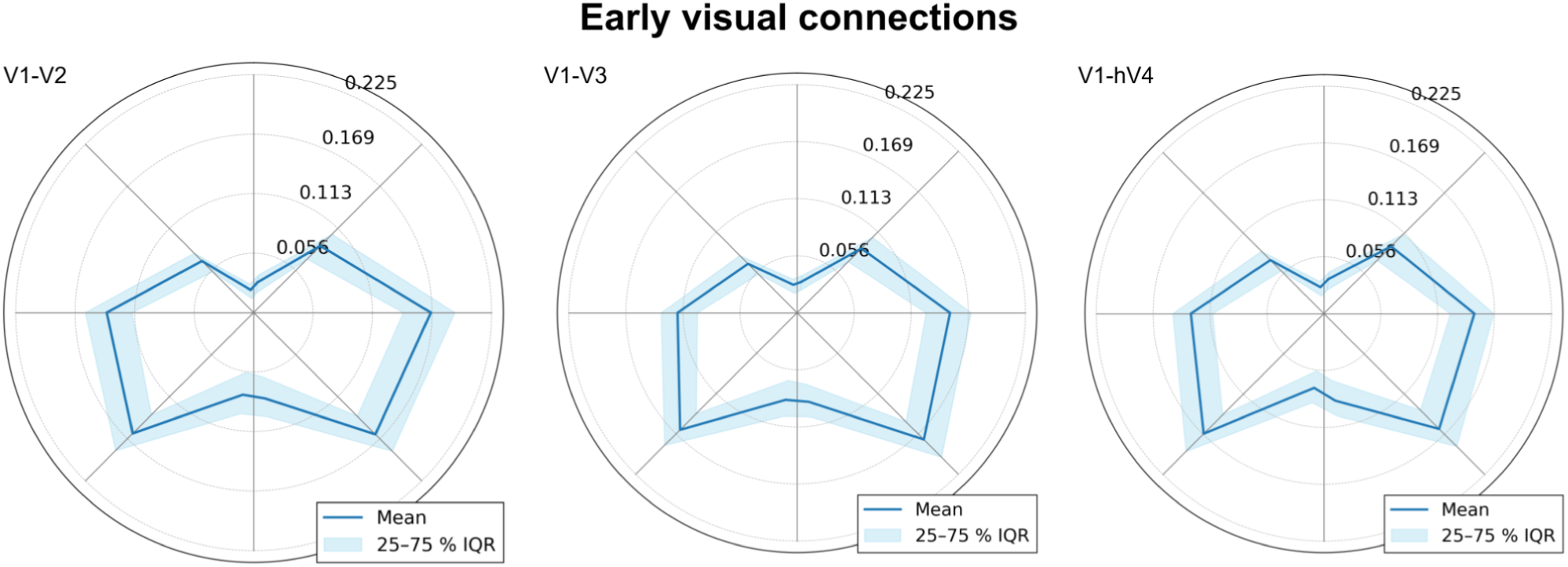
Visual meridian asymmetries across early visual connections. Polar-angle–resolved streamline density profiles for early visual pathways. Radar plots show mean streamline density (blue line) and interquartile range (25–75%, shaded area) across the four principal visual meridians: horizontal meridian (HM), vertical meridian (VM), lower vertical meridian (LVM), and upper vertical meridian (UVM). Connectivity between V1–V2, V1–V3, and V1–hV4 exhibits consistent visual meridian asymmetries, with higher streamline density along the horizontal relative to the vertical meridians (horizontal–vertical asymmetry, HVA) and stronger connectivity along the lower relative to the upper vertical meridian (vertical meridian asymmetry, VMA). These patterns indicate that structural connectivity within the early visual hierarchy reflects known perceptual asymmetries across the visual field.

For each pathway, the central tendency (mean) and variability (interquartile range, 25–75%) were computed, providing a compact representation of the distribution of connectivity strength across the visual field. Across multiple early visual pathways, including V1–V2, V1–V3, and V1–hV4, these profiles revealed consistent and systematic asymmetries in retinotopic connectivity. Specifically, streamline density was higher along the horizontal meridian than the vertical meridian, consistent with a horizontal–vertical asymmetry (HVA), and stronger along the lower than the upper vertical meridian, consistent with a vertical meridian asymmetry (VMA). These patterns mirror well-established perceptual asymmetries in human vision, in which visual performance is enhanced along the horizontal compared to the vertical meridian and in the lower than the upper vertical meridian^15–18^. Together, these results demonstrate that structural connectivity within the early visual cortex varies systematically with polar angle and reflects the spatial organization of perceptual performance across the visual field.

### Supplementary Results 5. Retinotopic connectivity asymmetry across higher-order visual areas

We next examined whether the retinotopically structured connectivity patterns observed in the early visual cortex extend to higher-order visual areas. Using VISCONN, we quantified streamline density profiles for connections involving higher-order regions, including hV4, LO1, LO2, TO1, and TO2.

Connectivity profiles were visualized using radial (radar) plots (see **Supplementary Fig. 6**) summarizing mean streamline density and interquartile range (25–75%) across the four principal visual meridians (HM, VM, LVM, UVM), as defined from retinotopic maps derived using population receptive field (pRF) modeling^3–5^. This representation enables direct comparison of how connectivity varies as a function of polar angle across different stages of the visual hierarchy. In contrast to early visual pathways, higher-order visual areas exhibited more heterogeneous and less spatially structured connectivity profiles. In several regions, the horizontal–vertical asymmetry (HVA), characterized by stronger connectivity along the horizontal relative to the vertical meridian, was absent. By comparison, the vertical meridian asymmetry (VMA), reflecting stronger connectivity along the lower relative to the upper vertical meridian, remained detectable in these higher-order areas.

These results indicate that the strong retinotopic organization of structural connectivity observed in the early visual cortex becomes progressively less pronounced along the visual hierarchy. The reduced expression of HVA and the more variable connectivity patterns across higher-order regions are consistent with increasing receptive field size and greater integration of information across the visual field in downstream visual areas^19,20^. Together, these findings suggest that while the early visual cortex preserves a tightly retinotopically organized connectivity structure, higher-order visual areas exhibit more heterogeneous connectivity profiles, reflecting a transition from spatially specific to more integrative representations of visual information.

**Supplementary Figure 6.**
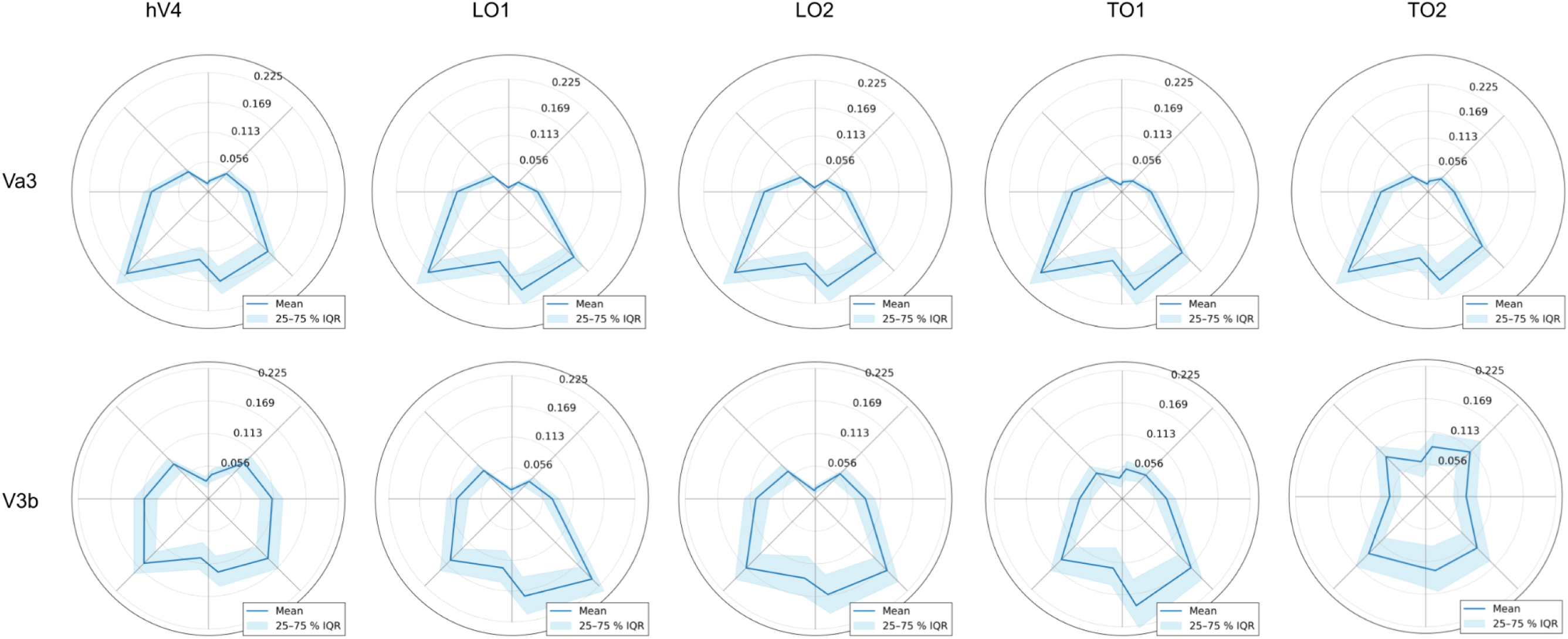
Visual meridian connectivity profiles across higher-order visual areas. Polar-angle-resolved streamline density profiles connections involving higher-order visual area, including hV4, LO1, LO2, TO1, TO2- V3a and V3b. Radar plots show mean streamline density (blue line) and interquartile range (25-75% shaded area) across the four Principal visual merdians. Compared with early visual pathways, higher-order regions exhibit more heterogeneous connectivity profiles. In many of These areas, the horizontal vertical asymmetry (HVA) is reduced or absent, whereas the vertical meridian asymmetry (VMA), characterized by stronger connectivity along the lower relative to the upper vertical meridian, remains detectable in several regions.

### Supplementary Results 6. Testing for Gyral Bias and Meridian Asymmetries

Diffusion MRI tractography is known to exhibit systematic biases related to cortical geometry, including a tendency for streamlines to preferentially terminate on gyral crowns relative to sulcal banks (gyral bias^21–23^).

To assess whether such biases could account for the observed meridian-dependent connectivity patterns, we evaluated the relationship between cortical geometry and streamline density across visual meridians.

Specifically, we tested whether differences in gyral versus sulcal sampling across retinotopic regions could influence estimates of horizontal–vertical (HVA) and vertical meridian (VMA) asymmetries. Importantly, cortical folding in early visual cortex is systematically related to retinotopic organization: vertical meridian representations tend to align with gyral crowns, whereas horizontal meridian representations are typically located within sulcal fundi^24^. Consistent with these observations, we replicated this meridian-specific relationship between cortical curvature and retinotopic organization across all datasets analyzed (HCP, Cam-CAN, and PING), confirming that the alignment between visual field representation and cortical geometry is robust across independent cohorts. This structure–function relationship implies that gyral bias would preferentially enhance connectivity estimates for vertical meridian representations relative to horizontal meridian representations.

Our analysis revealed that a gyral bias was present across most meridians, except for the horizontal meridian (see **Supplementary Fig. 7**). Notably, the magnitude of this bias was greater for vertical meridian representations than for horizontal meridian representations. Critically, this pattern predicts the opposite effect of the observed connectivity asymmetries: if gyral bias were driving the results, connectivity would be expected to be higher along the vertical meridian relative to the horizontal meridian. In contrast, our results consistently show higher connectivity along the horizontal meridian (see **Supplementary Fig. 8 and Fig. 3**). These findings indicate that the observed meridian asymmetries cannot be explained by tractography-related gyral bias; if anything, this bias would minimize the HVA we report here.

Together, these analyses demonstrate that the retinotopic connectivity asymmetries reported here are not attributable to known biases in tractography and instead reflect genuine spatial organization of structural connectivity within the visual system.

**Supplementary Figure 7.**
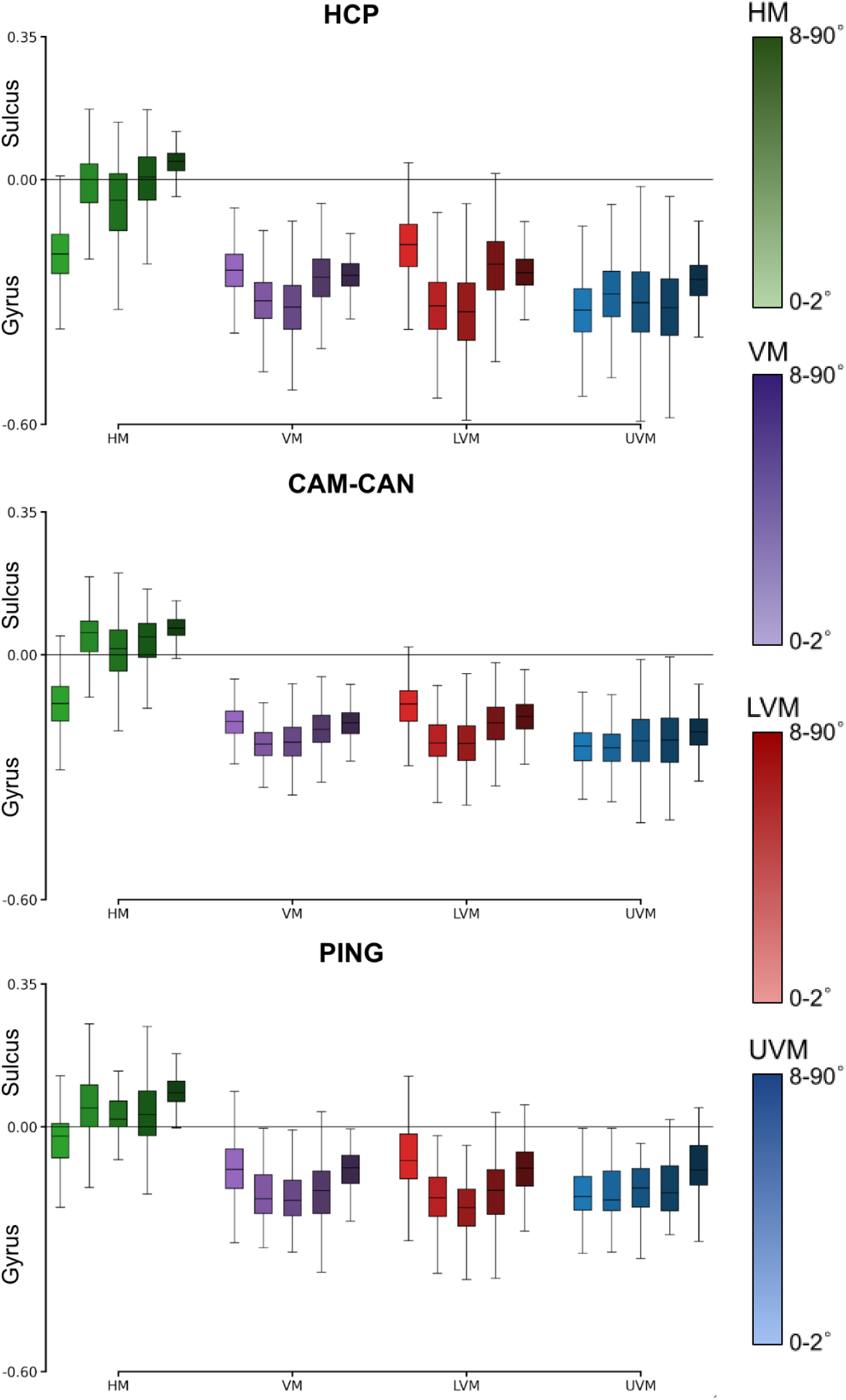
Gyral bias in visual field meridians across datasets. Boxplots show the distribution of mean cortical curvature values for each meridian and eccentricity bin in V1. Data are shown separately for the Human Connectome Project (HCP), Cambridge Centre for Ageing and Neuroscience (Cam-CAN), and Pediatric Imaging, Neurocognition, and Genetics (PING) datasets. Curvature values are derived from FreeSurfer surface reconstructions (negative values indicate gyri; positive values indicate sulci). Across datasets, most meridians exhibit a consistent gyral bias (i.e., negative curvature values), whereas the horizontal meridian shows reduced or absent bias. The magnitude of the gyral bias is greater for vertical meridian representations compared to horizontal meridian representations.

**Supplementary Figure 8.**
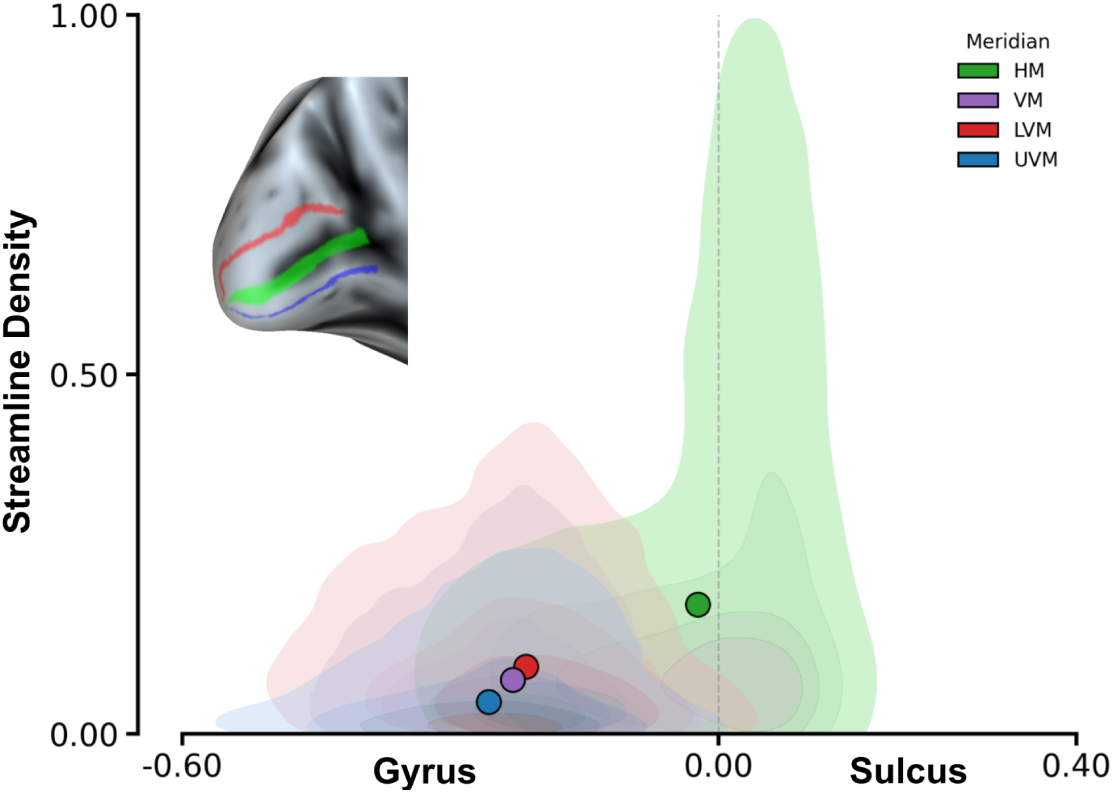
Distribution of Streamline Density Across Principal Visual Meridians as a Function of Cortical Curvature. Bivariate kernel density estimation (KDE) was used to characterize the relationship between streamline density and mean cortical curvature using data pooled across over 1,700 participants from the HCP, CAN-CAN, and PING datasets. Structural connectivity within primary visual cortex (V1) is shown across the cortical folding landscape for the principal visual field meridians, color-coded as horizontal (HM, green), vertical (VM, purple), lower vertical (LVM, red), and upper vertical (UVM, blue). Cortical curvature values were derived from surface reconstructions, with negative values (< 0) corresponding to gyral crowns and positive values (> 0) to sulcal fundi. Large filled circles denote group-level centroids for each meridian, summarizing the central tendency of the curvature–connectivity relationship across eccentricity bins and participants. Density contours (five levels) indicate that vertical meridian representations (VM; LVM and UVM) are predominantly localized within gyral cortex, consistent with their expected anatomical distribution, whereas horizontal meridian (HM) representations are shifted toward more positive curvature values, reflecting preferential localization within sulcal cortex. Notably, the highest streamline density values occur within the sulcal range and are primarily associated with the HM distribution.

## Supplementary Tables

**Supplementary Table 1.** Computational provenance of the VISCONN framework and associated analyses. Comprehensive list of the brainlife.io Apps, associated Digital Object Identifiers (DOIs), and GitHub source code repositories used in this study. The DOIs reference specific, versioned instances of each service executed on the brainlife.io platform, ensuring transparency and full reproducibility of the computational workflow.

|  |  |  |
| --- | --- | --- |
| Name | Brainlife DOI | Github Repository |
| FSL Anat (T1) | [10.25663/brainlife.app.273](https://doi.org/10.25663/brainlife.app.273) | [brainlife/app-fsl-anat](https://github.com/brainlife/app-fsl-anat) |
| Align T1 to ACPC Plane (HCP-based) | [10.25663/brainlife.app.800](https://doi.org/10.25663/brainlife.app.800) | [brainlife/app-hcp-acpc-alignment](https://github.com/brainlife/app-hcp-acpc-alignment) |
| FSL Anat (T2) | [10.25663/brainlife.app.350](https://doi.org/10.25663/brainlife.app.350) | [brainlife/app-fsl-anat](https://github.com/brainlife/app-fsl-anat) |
| Align T2 to ACPC Plane (HCP-based) | [10.25663/brainlife.app.116](https://doi.org/10.25663/brainlife.app.116) | [brainlife/app-hcp-acpc-alignment/tree/1.4](https://github.com/brainlife/app-hcp-acpc-alignment/tree/1.4) |
| Freesurfer | [10.25663/bl.app.0](https://doi.org/10.25663/bl.app.0) | [brainlife/app-freesurfer](https://github.com/brainlife/app-freesurfer) |
| Segment Thalamic Nuclei | [10.25663/brainlife.app.222](https://doi.org/10.25663/brainlife.app.222) | [brainlife/app-segment-thalamic-nuclei](https://github.com/brainlife/app-segment-thalamic-nuclei) |
| Tissue-type segmentation | [10.25663/brainlife.app.239](https://doi.org/10.25663/brainlife.app.239) | [brainlife/app-mrtrix3-5tt](https://github.com/brainlife/app-mrtrix3-5tt) |
| pRFs / Benson14-Retinotopy | [10.25663/brainlife.app.559](https://doi.org/10.25663/brainlife.app.559) | [brainlife/app-benson14-retinotopy/tree/v1.1](https://github.com/brainlife/app-benson14-retinotopy/tree/v1.1) |
| mrtrix3 preprocess | [10.25663/bl.app.68](https://doi.org/10.25663/bl.app.68) | [brainlife/app-mrtrix3-preproc/tree/1.7](https://github.com/brainlife/app-mrtrix3-preproc/tree/1.7) |
| FSL Brain Extraction (BET) on DWI | [10.25663/brainlife.app.163](https://doi.org/10.25663/brainlife.app.163) | [brainlife/app-FSLBET](https://github.com/brainlife/app-FSLBET) |
| Fit Constrained Deconvolution Model for Tracking | [10.25663/brainlife.app.238](https://doi.org/10.25663/brainlife.app.238) | [bacaron/app-mrtrix3-act](https://github.com/bacaron/app-mrtrix3-act) |
| Anatomically Constrained Tractography using precomputed 5tt & CSD | [10.25663/brainlife.app.297](https://doi.org/10.25663/brainlife.app.297) | [bacaron/app-mrtrix3-act](https://github.com/bacaron/app-mrtrix3-act) |
| mrtrix3 - WMC Anatomically Constrained Tractography (ACT) | [10.25663/brainlife.app.319](https://doi.org/10.25663/brainlife.app.319) | [brainlife/app-mrtrix3-act](https://github.com/brainlife/app-mrtrix3-act) |
| T1w-to-T1w Registration | [10.25663/brainlife.app.861](https://doi.org/10.25663/brainlife.app.861) | [gamorosino/app-registration-t1w/tree/v0.1](https://github.com/gamorosino/app-registration-t1w/tree/v0.1) |
| Apply Transformations to Tractography Data | [10.25663/brainlife.app.902](https://doi.org/10.25663/brainlife.app.902) | [gamorosino/app-tract-transform/tree/main](https://github.com/gamorosino/app-tract-transform/tree/main) |
| Structural network connectomics of visual white matter by polar angle and eccentricity | [10.25663/brainlife.app.784](https://doi.org/10.25663/brainlife.app.784) | [brainlife/app-visual-white-matter/tree/v1.2](https://github.com/brainlife/app-visual-white-matter/tree/v1.2) |
| Structural network connectomics of visual white matter by polar angle and eccentricity - Single Endpoints | [10.25663/brainlife.app.785](https://doi.org/10.25663/brainlife.app.785) | [brainlife/app-visual-white-matter/tree/v1.2](https://github.com/brainlife/app-visual-white-matter/tree/v1.2) |
| Tract Profile Computation | [10.25663/brainlife.app.911](https://doi.org/10.25663/brainlife.app.911) | [gamorosino/app-compute-tract-profile/tree/v0.1](https://github.com/gamorosino/app-compute-tract-profile/tree/v0.1) |
| Retinotopic Connectivity Matrix Computation | [10.25663/brainlife.app.910](https://doi.org/10.25663/brainlife.app.910) | [gamorosino/app-retinotopic-connectivity](https://github.com/gamorosino/app-retinotopic-connectivity) |
| Gyral Bias Analysis | [10.25663/brainlife.app.914](https://doi.org/10.25663/brainlife.app.914) | [gamorosino/app-gyral-bias-analysis](https://github.com/gamorosino/app-gyral-bias-analysis) |
| Fiber Bundle Skeleton Computation | [10.25663/brainlife.app.915](https://doi.org/10.25663/brainlife.app.915) | [gamorosino/app-bundle-skeleton](https://github.com/gamorosino/app-bundle-skeleton) |
| Streamline Cleaning | [10.25663/brainlife.app.912](https://doi.org/10.25663/brainlife.app.912) | [gamorosino/app-streamline-cleaning/tree/v0.1](https://github.com/gamorosino/app-streamline-cleaning/tree/v0.1) |
| Tractogram clustering | [10.25663/brainlife.app.916](https://doi.org/10.25663/brainlife.app.916) | [gamorosino/app-tract-mbkm-cluster](https://github.com/gamorosino/app-tract-mbkm-cluster) |
| Apply Transformations to Tractography Data | [10.25663/brainlife.app.902](https://doi.org/10.25663/brainlife.app.902) | [gamorosino/app-tract-transform/tree/v2.0](https://github.com/gamorosino/app-tract-transform/tree/v2.0) |
| Map Retinotopic Connections | [10.25663/brainlife.app.913](https://doi.org/10.25663/brainlife.app.913) | [gamorosino/app-map-retinotopic-tracts/tree/main](https://github.com/gamorosino/app-map-retinotopic-tracts/tree/main) |
| Vertical Occipital Fasciculus (VOF) Coverage Quantification | [10.25663/brainlife.app.917](https://doi.org/10.25663/brainlife.app.917) | [gamorosino/app-vof-coverage](https://github.com/gamorosino/app-vof-coverage) |

**Supplementary Table 2.**
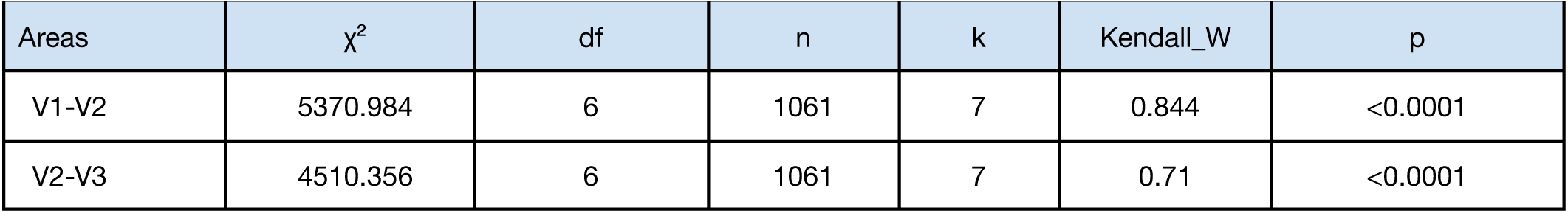
One-way repeated measures analysis of variance by ranks (Friedman’s ANOVA) for like-to-like connectivity across visual areas. A non-parametric one-way repeated-measures analysis (Friedman test) was used to assess differences in streamline density across retinotopic distance shells (0 to ≥6 DVA) for V1–V2 and V2–V3 connectivity. Reported values include the Friedman test statistic (χ²), degrees of freedom (df), number of participants (n), number of conditions (k), and Kendall’s W as a measure of effect size. Across both area pairs, a significant effect of retinotopic distance was observed (all p<0.001), with large effect sizes (Kendall’s W = 0.844 for V1–V2 and 0.71 for V2–V3).

| Areas | $\chi^2$ | df | n | k | Kendall_W | p |
| --- | --- | --- | --- | --- | --- | --- |
| V1-V2 | 5370.984 | 6 | 1061 | 7 | 0.844 | <0.0001 |
| V2-V3 | 4510.356 | 6 | 1061 | 7 | 0.71 | <0.0001 |

**Supplementary Table 3.** Post hoc Wilcoxon signed-rank tests for the V1–V2 like-to-like analysis. Pairwise comparisons tested whether streamline density in the main diagonal (same DVA; iso-eccentric connections) was greater than in successive off-diagonals representing increasing eccentricity difference (±1 to ≥6 DVA). Reported values include the Wilcoxon signed-rank statistic (W), its normal-approximation Z value, the associated effect sizes (r and paired-samples Cohen’s d), and Holm-corrected p-values. Across all comparisons, connectivity in the same-DVA shell was significantly greater than in all off-diagonal shells, with effect sizes increasing monotonically as a function of retinotopic separation.

| Comparison | Direction | W | Z | r | d | p |
| --- | --- | --- | --- | --- | --- | --- |
| same DVA vs $\pm 1$ DVA | greater | 489803 | 20.928 | 0.643 | 0.711 | <0.0001 |
| same DVA vs $\pm 2$ DVA | greater | 544074 | 26.371 | 0.81 | 1.009 | <0.0001 |
| same DVA vs $\pm 3$ DVA | greater | 553084 | 27.275 | 0.838 | 1.132 | <0.0001 |
| same DVA vs $\pm 4$ DVA | greater | 559819 | 27.951 | 0.858 | 1.25 | <0.0001 |
| same DVA vs $\pm 5$ DVA | greater | 561831 | 28.152 | 0.865 | 1.322 | <0.0001 |
| same DVA vs $> 6$ DVA | greater | 562277 | 28.197 | 0.866 | 1.405 | <0.0001 |

**Supplementary Table 4.** Post hoc Wilcoxon signed-rank tests for the V2–V3 like-to-like analysis. Pairwise comparisons tested whether streamline density in the main diagonal (same DVA; iso-eccentric connections) was greater than in successive off-diagonals representing increasing eccentricity difference (±1 to ≥6 DVA). Reported values include the Wilcoxon signed-rank statistic (W), its normal-approximation Z value, the associated effect sizes (r and paired-samples Cohen’s d), and Holm-corrected p-values. Across all comparisons, connectivity in the same-DVA shell was significantly greater than in all off-diagonal shells, with effect sizes increasing monotonically as a function of retinotopic separation.

| Comparison | Direction | W | Z | r | d | p |
| --- | --- | --- | --- | --- | --- | --- |
| same DVA vs $\pm 1$ DVA | greater | 500532 | 22.088 | 0.679 | 0.778 | <0.0001 |
| same DVA vs $\pm 2$ DVA | greater | 525833 | 24.63 | 0.757 | 0.905 | <0.0001 |
| same DVA vs $\pm 3$ DVA | greater | 550932 | 27.151 | 0.834 | 1.122 | <0.0001 |
| same DVA vs $\pm 4$ DVA | greater | 558045 | 27.865 | 0.856 | 1.286 | <0.0001 |
| same DVA vs $\pm 5$ DVA | greater | 560214 | 28.083 | 0.863 | 1.394 | <0.0001 |
| same DVA vs $> 6$ DVA | greater | 500532 | 22.088 | 0.679 | 0.778 | <0.0001 |

**Supplementary Table 5.** One-way repeated measures analysis of variance by ranks (Friedman’s ANOVA) for meridian asymmetries in V1-V2 structural connectivity across datasets. A non-parametric one-way repeated-measures analysis (Friedman’s ANOVA) was used to assess differences in streamline density across visual field meridians (HM, LVM, and UVM) in the HCP, CAM-CAN, and PING datasets. Reported values include the Friedman test statistic (χ²), degrees of freedom (df), number of participants (n), number of conditions (k), and Kendall’s W as a measure of effect size. Across all datasets, a significant effect of visual meridian was observed (all p < 0.0001), with large effect sizes (Kendall’s W ranging from 0.765 to 0.956), indicating a robust and consistent modulation of structural connectivity by retinotopic location.

| Dataset | $\chi^2$ | df | n | k | Kendall_W | p |
| --- | --- | --- | --- | --- | --- | --- |
| HCP | 2026.36 | 2 | 1061 | 3 | 0.956 | <0.0001 |
| CAM-CAN | 893.866 | 2 | 584 | 3 | 0.765 | <0.0001 |
| PING | 195.528 | 2 | 106 | 3 | 0.922 | <0.0001 |

**Supplementary Table 6.** Post hoc Wilcoxon signed-rank tests for meridian asymmetries in V1-V2 structural connectivity across datasets. Pairwise comparisons tested whether streamline density differed between visual field meridians, specifically assessing horizontal–vertical asymmetry (HVA; HM > VM) and vertical meridian asymmetry (VMA; LVM > UVM) in the Human Connectome Project (HCP), PING, and CAM-CAN datasets. Reported values include the Wilcoxon signed-rank statistic (W), its normal-approximation Z value, and associated effect sizes (r and paired-samples Cohen’s d), along with Holm-corrected p-values. Across datasets, a robust and consistent VMA (LVM > UVM) was observed, with large effect sizes in all cohorts. In contrast, the HVA (HM > VM) was significant in the HCP and PING datasets but not in CAM-CAN, where effect sizes were near zero, and the direction of the effect was not consistent across participants.

| Dataset | Comparison | Direction | W | Z | r | d | p |
| --- | --- | --- | --- | --- | --- | --- | --- |
| HCP | HM vs VM | greater | 562070 | 28.176 | 0.865 | 2.513 | <0.0001 |
|  | LVM UVM | greater | 560882 | 28.057 | 0.862 | 1.467 | <0.0001 |
| CAM-CAN | HM vs VM | greater | 166625 | 19.909 | 0.824 | 1.382 | <0.0001 |
|  | LVM UVM | greater | 158643 | 17.952 | 0.743 | 0.964 | <0.0001 |
| PING | HM vs VM | greater | 5671 | 8.937 | 0.868 | 2.315 | <0.0001 |
|  | LVM UVM | greater | 5091 | 7.109 | 0.69 | 0.885 | <0.0001 |

